# Pneumococcal within-host diversity during colonisation, transmission and treatment

**DOI:** 10.1101/2022.02.20.480002

**Authors:** Gerry Tonkin-Hill, Clare Ling, Chrispin Chaguza, Susannah J Salter, Pattaraporn Hinfonthong, Elissavet Nikolaou, Natalie Tate, Andrzej Pastusiak, Claudia Turner, Claire Chewapreecha, Simon DW Frost, Jukka Corander, Nicholas J Croucher, Paul Turner, Stephen D Bentley

## Abstract

Characterising the genetic diversity of pathogens within the host promises to greatly improve surveillance and reconstruction of transmission chains. For bacteria, it also informs our understanding of inter-strain competition, and how this shapes the distribution of resistant and sensitive bacteria. Here we study the genetic diversity of *Streptococcus pneumoniae* within individual infants and their mothers by deep sequencing whole pneumococcal populations from longitudinal nasopharyngeal samples. We demonstrate deep sequencing has unsurpassed sensitivity for detecting multiple colonisation, doubling the rate at which highly invasive serotype 1 bacteria were detected in carriage compared to gold-standard methods. The greater resolution identified an elevated rate of transmission from mothers to their children in the first year of the child’s life. Comprehensive treatment data demonstrated infants were at an elevated risk of both the acquisition, and persistent colonisation, of a multidrug resistant bacterium following antimicrobial treatment. Some alleles were enriched after antimicrobial treatment, suggesting they aided persistence, but generally purifying selection dominated within-host evolution. Rates of co-colonisation imply that in the absence of treatment, susceptible lineages outcompeted resistant lineages within the host. These results demonstrate the many benefits of deep sequencing for the genomic surveillance of bacterial pathogens.

## Background

*Streptococcus pneumoniae* is a highly recombinogenic human nasopharyngeal commensal and respiratory pathogen that causes high rates of pneumonia, bacteremia and meningitis, particularly in young children and the elderly (1, 2). Individual strains have been observed to diversify through point mutation, recombination and mobile element acquisition during nasopharyngeal carriage and disease, affecting antimicrobial resistance, susceptibility to vaccine-induced immunity, and the inference of transmission networks (3, 4). Further complexity arises from simultaneous carriage of multiple strains. The co-existence of resistant and sensitive strains, and the re-structuring of populations following vaccine introduction, both suggest that within-host competition between strains could play a major role in the population dynamics of *S. pneumoniae* (5–7).

As with many bacterial pathogens, surveillance of *S. pneumoniae* has been revolutionised by large-scale whole genome sequencing (WGS) efforts, which have greatly enhanced our ability to track antibiotic resistant and vaccine evading lineages at the population level (8–11). However, similar to other bacterial pathogens, genomic surveillance of *S. pneumoniae* has typically relied on the analysis of a representative genome generated from a single colony from a patient or carrier (9, 12). This limits the sensitivity of surveillance, as carriage of multiple distinct pneumococcal lineages is frequent in areas with high prevalence. Infant pneumococcal carriage rates can be as high as 100% in parts of Africa and Southeast Asia (13–16).

Previous studies of within-host diversity in bacteria have mostly relied on separately sequencing the genomes of multiple purified colonies, isolated from a single person, which incurs a significant time and financial cost (17, 18). Conversely, within-host population deep sequencing (PDS) involves sequencing a pool of hundreds of colonies from a sample producing a high depth sampling of within-host diversity. While this has been shown to provide a more detailed picture of the genetic diversity within the host (19), these analyses have predominantly focused on laboratory studies (20), relatively small outbreaks (21) or clinical isolates taken from symptomatic patients, particularly for bacterial species known to colonise patients with cystic fibrosis or other chronic lung diseases (22, 23).

Here, using a deep sequencing approach, we study the evolutionary dynamics of *S. pneumoniae* within healthy carriers, and during episodes of illness and antibiotic treatment, while also examining the potential utility of within-host population sequencing in surveillance. We analyse data from nearly 4000 samples collected during a large longitudinal carriage study conducted between 2007-2010 in the Maela refugee camp on the border of Thailand and Myanmar. Nasopharyngeal swabs were collected from 965 infants, and a subset of their mothers, from birth until 24 months of age (Figure 1A).

**Fig. 1.**
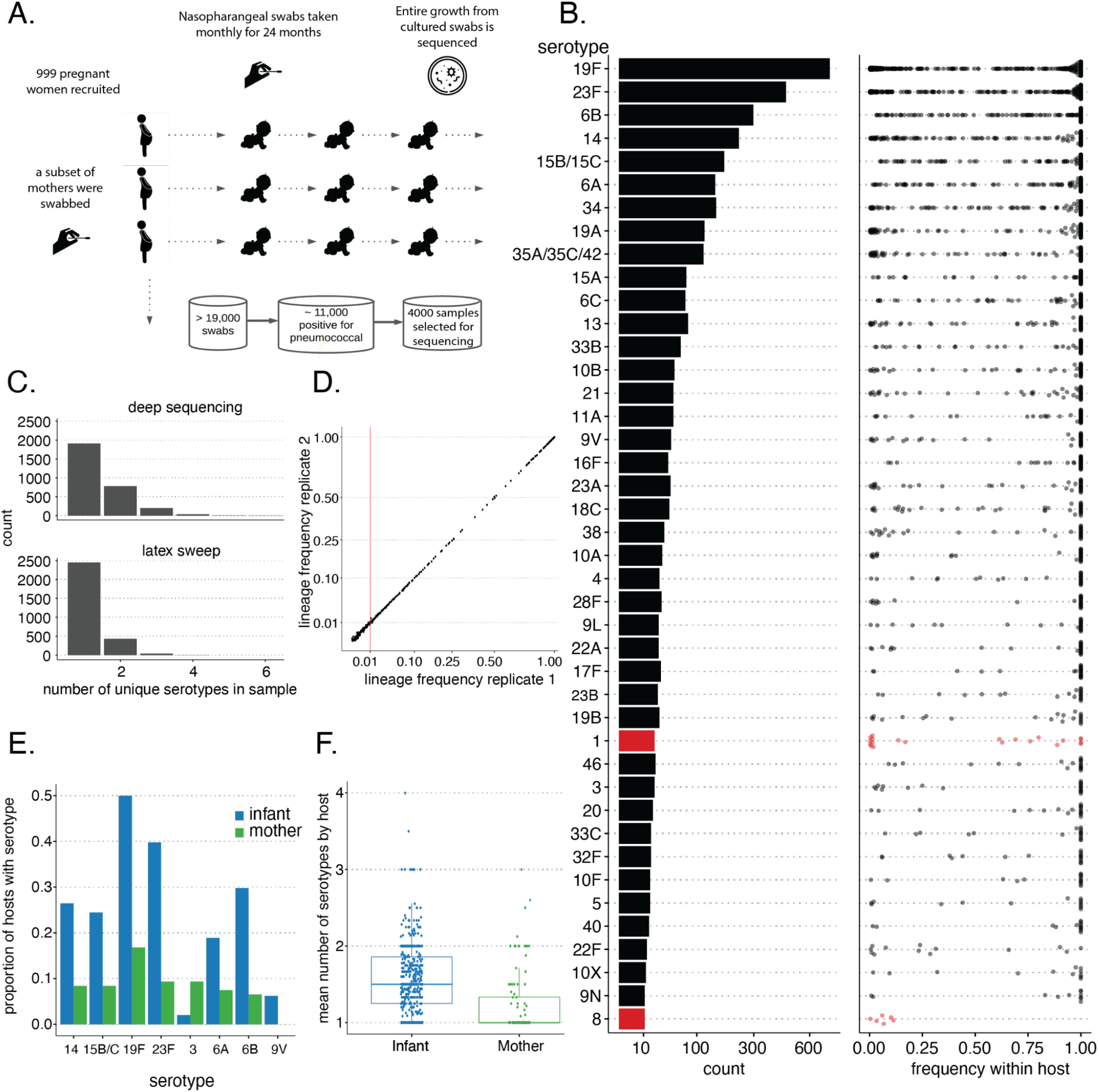
a.) A schematic of the sampling study design. b.) A bar plot indicating the number of times each serotype was observed across all deep sequenced samples. The distribution of the corresponding within-host frequencies of these serotypes is given in the adjacent plot with overlapping points separated to indicate the density at each position along the x-axis. Lineages with ambiguous serotype calls were excluded from this plot. c.) Histograms indicating the distribution of the number of unique serotypes observed using either PDS or latex sweeps. d.) Comparisons between the estimated GPSC lineage frequencies in 192 samples which were sequenced in replicate. The vertical red bar indicates the minimum frequency required for consideration in the mSWEEP pipeline. e.) Barplots indicating the differences in the representation of serotypes between mothers and infants. f.) Distribution in the mean number of serotypes observed in mothers and their infants.

### Within-host population deep sequencing provides accurate lineage and serotype calls

We first examined whether accurate lineage and serotype calls could be made from the pooled data obtained from deep sequencing hundreds of colonies from plate scrapes of pneumococci grown on selective agar referred to as population deep sequencing (PDS). Lineages were defined using the Global Pneumococcal Sequencing Cluster (GPSC) nomenclature (9). We used a dual approach of both deconvoluting the mixed samples and running standard analyses, as well as using methods designed for analysing population sequencing data directly (Methods). We calibrated and verified the approach using a subset of 1158 samples for which single colonies had been selected and sequenced in a previous study (24). In addition, we made use of 44 artificial laboratory mixtures for which sequencing data was also available (25). To check for potential processing artefacts, 192 of the selected samples were sequenced in replicate with separate PCR amplification and library preparation steps. The culture step was also replicated in a further 8 samples.

The within-host PDS approach reliably detected lineages (GPSCs) within each sample, with a precision and recall of 100% and 93% respectively on the artificial laboratory samples. It correctly re-identified 97.1% (1149/1158) of the lineages present in the larger set of carriage isolates (Supplementary Figure 1). For the culture and sequencing replicates that passed initial QC, the approach achieved an accuracy of 100% (3/3) and 97.5% (157/161). Figure 1D, shows that the estimated frequency of each lineage was highly concordant between sequencing replicates with a correlation of > 0.99 (*p* < 1*e* 3, Fisher’s Z-transform). Although lower, the concordance observed within the three culture replicates (*ρ* = 0.94, *p* = 0.059) was still strong. This indicates that the estimated frequencies are robust to potential artefacts of the experimental pipeline, allowing us to confidently interpret relative changes in frequencies.

### Population deep sequencing reveals hidden diversity

Using PDS we identified 23.6% (813/3450) more serotypes compared to the most common method of identifying multiple colonisation (latex sweep) (14) (Figure 1C). Due to difficulties in distinguishing ambiguous or poor quality serotype calls from non-typables we assigned such lineages with an ‘unknown’ serotype. Multiple distinct serotypes were observed in 1028/2940 (35%) of samples, further highlighting the substantial genetic diversity that is obscured by standard surveillance using single representative genomes. The increased sensitivity was supported by microarray data on a subset of 32 samples, which identified all 49 serotypes found by PDS, compared to the 32 found using latex sweeps (14). Unlike PDS, microarray data only indicates the presence and absence of known genes and serotypes, and does not provide data over the entire genome.

Rates of multiple colonisation were significantly higher in infants than in their mothers (p<1e-3, Welch two sample t-test) (Figure 1F). The most common serotypes, including 19F and 23F, were also significantly more likely to be found in infants (Figure 1E), which is consistent with a greater repertoire of adaptive immunity in adults (adjusted p < 0.05, Welch two sample t-test) (26). In agreement with past studies, serotype 3 was the only serotype to be found more frequently in mothers (Figure 1E) (27–30). It has been postulated that the high rate of invasive disease due to serotype 3 in adults may correlate with high antibody levels in children, which then wane (31–33).

Other “epidemic” serotypes (e.g. 1, 2, 5, 7F, 8 and 12F) are known for causing outbreaks of disease in adults, despite rarely being detected in infant carriage (34). Strikingly, we found such types were often present at low frequencies within the host (Figure 1B). In particular, serotypes 1 and 8, and the associated GPSCs 2 and 28, were found at lower frequencies than other types (adj p-value < 0.05, Kolgomorov-Smirnov test). In 11/20 (55%) of the observed cases of serotype 1 in our dataset, it was found as the minority serotype in multiple colonisation. This could partly explain its low detection rate in previous carriage studies (8, 9, 35) which typically only detect the dominant strain in each sample. Given invasiveness is usually calculated by comparing carriage and disease rates, this suggests that current estimates of the invasiveness of serotype 1 may be inflated. However, despite PDS identifying over twice as many serotype 1 lineages, the overall prevalence of this serotype was still low, making up <1% of all distinct serotype-host pairs in the dataset. Consequently, this serotype still appears to be highly invasive, justifying its targeting by current vaccines (36).

We found that PDS identified an additional 14.6% (520/3557) resistance elements, including known resistance SNPs and mobile genetic elements, when compared to using standard pipelines on the set of 1158 single colony whole genome sequences (Supplementary Figure 1C). This indicates that resistant lineages are frequently found alongside susceptible lineages within the same host. The rate of resistance in samples taken from infants was significantly higher than in mothers for 4/14 antibiotic classes, which corresponds with the difference in the composition of lineages observed between mothers and children (Supplementary Figure 3, adj p <0.05). Thus, routine PDS provides substantial improvements over alternative approaches in the surveillance of pneumococcal resistance, especially in children, where rates of multiple colonisation are higher.

### Inference of mother-infant transmission indicates an age-dependent rate of asymmetric transmission and an incomplete transmission bottleneck

Deep within-host population sequencing also allows for improved estimates of transmission links (3, 37). To provide a robust measure of the strength of a transmission link between any two samples in our dataset we adapted the TransCluster algorithm to account for within-host diversity information (38). As a substitute to the classic SNP distance commonly used in studies of transmission, for each pair of samples that shared a common GPSC we counted the number of sites along the corresponding reference genome for that GPSC where no matching allele could be found in both samples (Supplementary Figure 7). A Multinomial-Dirichlet distribution for the allele counts at each site in each sample was used to account for the variable sequencing depth across the genome (Methods). This approach provides a lower bound estimate of the genetic divergence separating any pair of pneumococcal genomes within each of the two samples, while allowing for the possibility of multiple colonisation.

There was a strong association between the probability of direct transmission, as inferred by the adapted TransCluster algorithm independently of location data, and the geographic proximity of households (Figure 2A-B). This association remained after excluding within-household pairs involving mothers and their children, suggesting that children living closer to detected cases of more invasive strains are at higher risk, which could motivate local interventions to reduce transmission in outbreaks. Of the inferred close transmission links (estimated to involve either 0 or 1 intermediate hosts), 80.9% (871/1077) contained at least one sample found to carry multiple pneumococcal lineages. Part of this can be attributed to the high level of multiple colonisation in the cohort, but this nevertheless suggests that any transmission inference study that neglects to account for multiple colonisation could substantially underestimate the number of close transmission links.

**Fig. 2.**
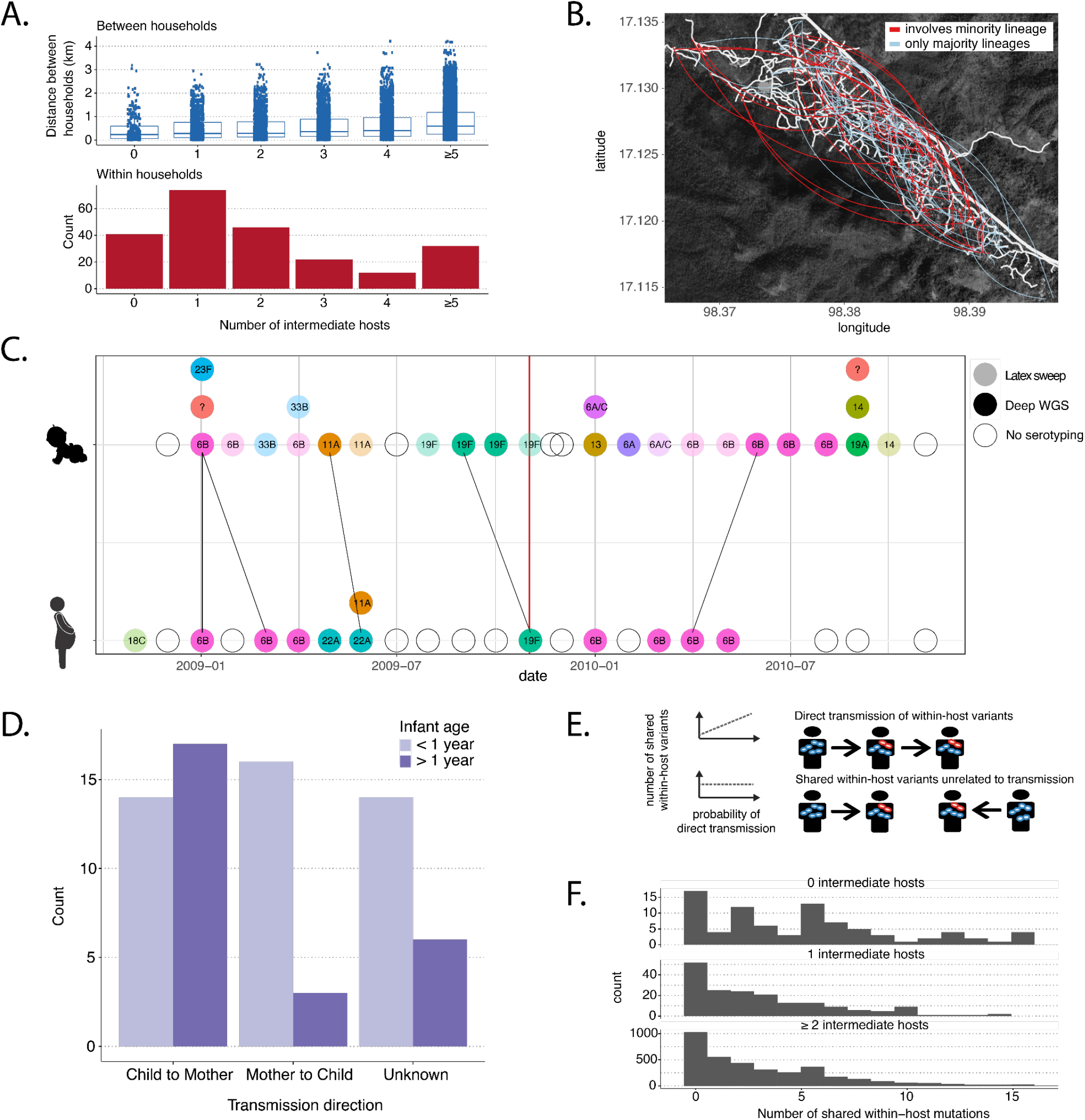
a.) The distribution of pairwise geographic distances between each household versus the number of intermediate transmission events as inferred using the modified TransCluster algorithm (above). The distribution of estimated intermediate transmission events within households (below). b.) A map of the Maela refugee camp with inferred direct transmission links overlaid. Blue lines indicate links that would typically be inferred using a representative genome per sample while red lines indicate additional links that were found using PDS. c.) A representative mother/child pair indicating how transmission direction was inferred. Circles indicate the serotypes present with PDS data available for those coloured in bold. Black lines indicate close transmission links inferred using the TransCluster algorithm with the red line indicating the time the child was one year old. d.) The distribution of the direction of transmission between mother and child split by whether the transmission event occurred before or after the child turned one. e.) A schematic demonstrating that we would expect to see an elevated rate of polymorphic sites amongst close transmission pairs. f.) The distribution of the number of shared polymorphic sites in transmission pairs involving 0, 1 or *≥* 2 intermediate hosts. The elevated number of variants involving 0 intermediates hosts indicates a mean bottleneck size *≥* 1.

This high resolution approach also allowed us to scrutinise the transmission bottleneck, which is the point of the pneumococcal lifecycle blocked by immunity induced by current vaccines (39). Laboratory experiments have suggested that there is likely to be a very tight bottleneck in the transmission of *S. pneumoniae*, consisting of as little as a single bacterial cell (40). To understand how well these experiments generalise to transmission in human hosts, we took a conservative approach, using only samples containing a single strain. We compared the distribution of the number of shared polymorphic sites in samples with the most likely number of intermediate hosts, as inferred using the TransCluster algorithm (Figure 2E). The effects of hypermutable sites, sequencing errors and multiple infections, which have been shown to confound efforts to estimate the size of the transmission bottleneck, are likely to be similar irrespective of how close two samples are in the transmission chain (41, 42). Thus, any increase in the number of shared polymorphic sites between samples that are likely to be related by recent transmission is likely to be the result of multiple genotypes being transmitted (Figure 2E). The significant increase in the number of shared polymorphic sites found in putative direct transmission pairs relative to those estimated to involve intermediate hosts suggests that while tight, the transmission bottleneck between the donor and recipient is likely greater than one (p<1e-3, Poisson regression) (Figure 2F). Mouse models of pneumococcal transmission have indicated that this bottleneck is likely to occur following exit but before establishment in the recipient host (40).

To examine transmission within the home, we next considered the 47 mother/child pairs for which a transmission link involving zero or one intermediate host was inferred using the TransCluster algorithm. To estimate a plausible direction of transmission we required that the infector must have acquired the relevant lineage prior to the infectee, and that there could be at most one negative or missing sample in the infector in the 2 months prior to the infectee becoming infected with the same lineage (Figure 2C). The vast majority (16/19) of mother to child transmissions occurred in the first year of the infant’s life (Figure 2D). This was significantly different from the child to mother transmissions (14/31, p=0.015 Chisquared test). This asymmetry is consistent with <1 year old infants being more susceptible to infections from within the household, and with the high proximity between mother and child. The exposure risk posed by adults has been observed in previous studies (43–45) with routine vaccination of older children found to not have a significant effect on vaccine type carriage rates in unvaccinated infants (46). Taken together this suggests there may be a benefit to a vaccination campaign targeting mothers or other adults with high contact rates to young infants before herd immunity in the adult population is established. However, this would not reduce the risk posed by non-vaccine type lineages.

### Within-host minority variants are characterised by strong purifying selection and a distinct mutational spectrum

To investigate selection acting at the scale of individual lineages within the host we restricted our analysis to within-host single nucleotide variants found in samples involving only a single pneumococcal lineage (Figure 3C). This avoided the potential for biases or errors to be introduced by the deconvolution of mixed samples. Minority variants were called using a conservative pipeline that included a scan statistic to filter out regions that were likely to be affected by homologous recombination, gene duplications and similarity with bacteriophages and other bacterial species (Methods). Many of the regions identified by this scan included genes coding for major pneumococcal autolysin proteins (including LytA) and other surface-associated choline binding proteins (CBP, including Pneumococcal Surface Proteins A and C, PspA and PspC) and the Tuf elongation factor (Supplementary Figure 6). Homologs to LytA and CBP domains are frequently found in pneumococcal phages or co-colonising bacterial species (47, 48). Phages may facilitate pneumococcal diversification and recombination in these regions is often enriched on the terminal branches of pneumococcal phylogenies as integration is often detrimental to the bacterium (48, 49).

**Fig. 3.**
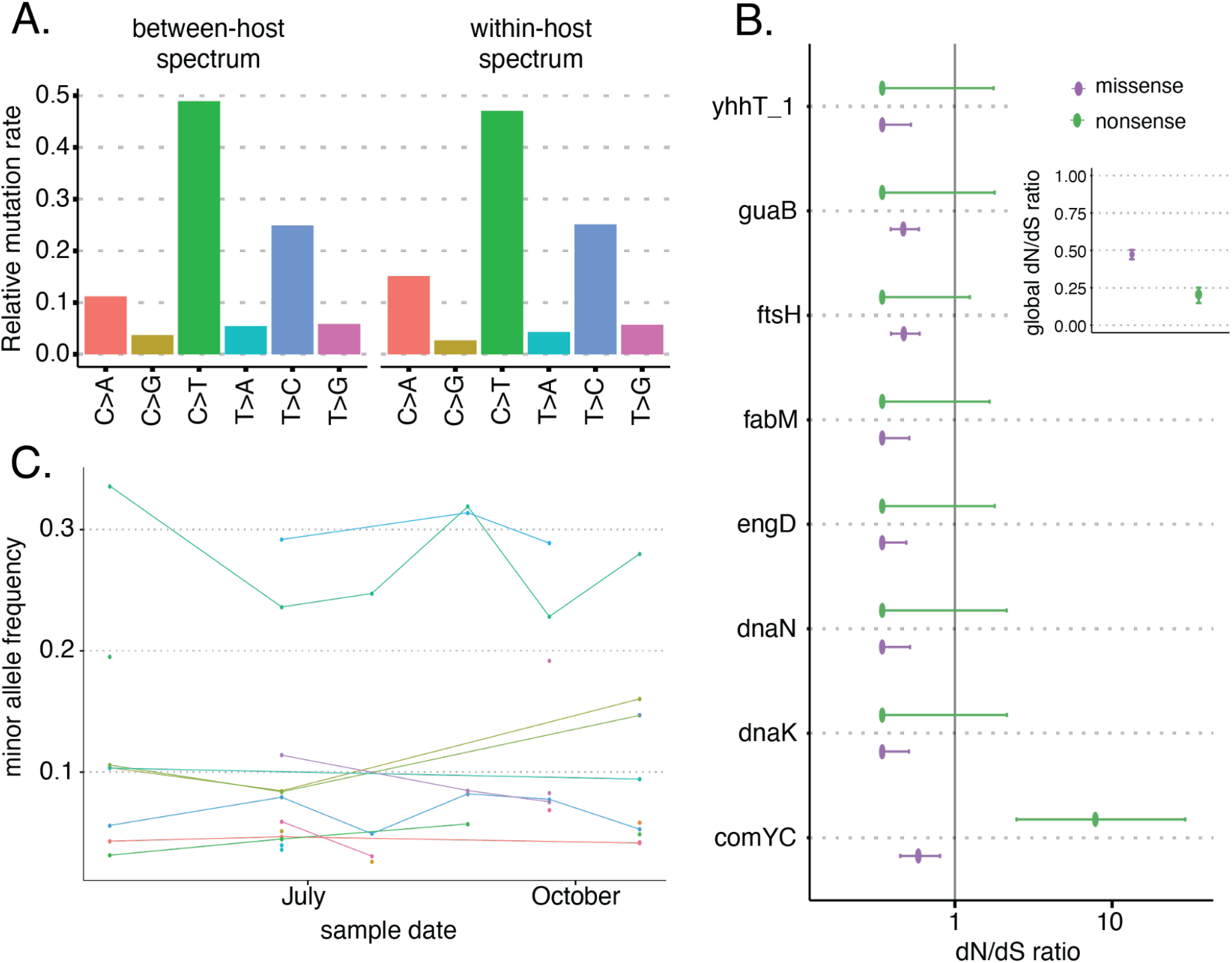
a.) The relative fraction of different single nucleotide base changes found within the host in samples involving only a single pneumococcal lineage compared to those changes observed between hosts inferred using ancestral state reconstruction. b.) dN/dS ratios for genes found to be under significant selection within the host. c.) An example of the within-host variant allele frequencies in consecutive samples of an infant carrying a single pneumococcal lineage.

The remaining within-host single nucleotide variants displayed a similar mutational spectrum to that found in the genome phylogenies constructed from single colonies taken from separate hosts (Figure 3A, Supplementary Figure 17) (24). This supports the quality of the calls and indicates that similar mutational processes are acting across the different timescales observed within hosts as opposed to between hosts. Although the overall spectra were similar, we observed an elevated number of C−>A transversions with weak sequence context in the deep-sequencing calls (p<1e-3, permutation test). This is consistent with the expected spectra of oxidative- and deamination-induced damage, which is typically reduced in frequency by purifying selection over longer time scales (50). A similar enrichment of C−>A mutation has also been seen in *E. coli* over short timescales within laboratory experiments, which has been hypothesised to be driven by the misincorporation of adenines into cytosine sites (51). Finally, pneumococci carry the *spxB* gene which secretes hydrogen peroxide and has been shown to cause DNA damage to host lung cells and may be a significant contributing factor to the observed within-host mutational spectrum of the bacterium itself (52).

To investigate signatures of selection within the single nucleotide variant calls we considered the dN/dS ratios calculated using a modified version of the dNdScv package (53). dNdScv uses a Poisson framework allowing for more complex substitution models that account for context dependence and the non-equilibrium of substitutions in estimating dN/dS ratios (53). This is particularly important in the case of sparse mutations in low recombination environments, as is the case in pneumococcal carriage over short timescales.

The resulting dN/dS ratios demonstrated clear purifying selection, particularly against nonsense mutations (Figure 3B). This is consistent with the strong selective pressures acting within the host with similar ratios observed in other respiratory pathogens (17, 23, 41, 54). Significant purifying selection was also observed at the level of individual genes, with only a single competence related gene (*comYC*) having an elevated rate of nonsense mutations (BH adjusted p-value < 0.05) (Figure 3B). This reflects the frequent insertion of pneumococcal prophage into *comYC*, causing premature stop codons that prevent the host cell from undergoing transformation, and have been associated with a reduced duration of carriage (55–57).

Those genes for which there was strongest evidence of purifying selection from dN/dS were frequently linked to the pneumococcal stress response including those for the heat shock proteins *dnaK* and *ftsH*, as well as *fabM,* which is necessary for survival in high acidity environments (58, 59). Purifying selection was also observed in the purine biosynthesis gene *guaB*. Multiple-antigen *S. pneumoniae* vaccines which include DnaK, as well as other heat shock proteins, have been shown to protect against lethal pneumococcal challenge (60). FabM has also been suggested as a potential target for novel chemotherapeutic agents (61). The observed purifying selection indicates that it may be difficult for the pneumococcus to adapt to treatments targeting these genes over short timescales. Although we are able to detect purifying selection, using dN/dS we did not find evidence for short term adaptive evolution in any genes. This likely reflects the long term commensal lifestyle of *S. pneumoniae* in contrast to that seen in environmental or immunocompromised patient pathogens (22, 23).

### Evidence of competition between resistant and susceptible lineages has important implications for pneumococcal population structure

The majority of multiple colonisation events between different GPSCs were observed at only a single time point 92.3% (712/771), indicating that long term multiple colonisation of the same lineages is rare. However, we did observe a number of carriage events where two lineages coexisted for well over the month-long time period between routine sampling. This suggests that competition between lineages within the host is not always strong enough for one to exclude the other (Supplementary Figure 14). Despite the large sample size, we did not have the statistical power to identify any preferential co-colonisation between particular pneumococcal lineages due to the very large number of possible combinations.

While resistant lineages were frequently observed to co-colonise with susceptible lineages, this occurred less frequently than would be expected given the background frequency of resistant lineages within the Maela camp (Figure 4A). Using a generalised linear mixed model to control for both host and pneumococcal population structure effects (Methods) we found that rates of resistance in multiple colonisation were significantly lower than expected in 5/14 antibiotic classes, including penicillin. This is consistent with resistant lineages being outcompeted by susceptible lineages within the host as a result of the fitness costs associated with resistance (62). Many models of the maintenance of antibiotic resistance in pneumococcal populations rely on assumptions about the competition between resistant and susceptible lineages within the host (5, 6, 63). However, currently most studies have relied upon serotype data alone to determine multiple colonisation rates, which do not indicate whether the underlying lineages are resistant to antibiotics. This result confirms that resistant and susceptible lineages are found to co-colonise the same host, and that the expected fitness costs of resistance observed in laboratory experiments are consistent with the population dynamics observed in natural pneumococcal carriage.

**Fig. 4.**
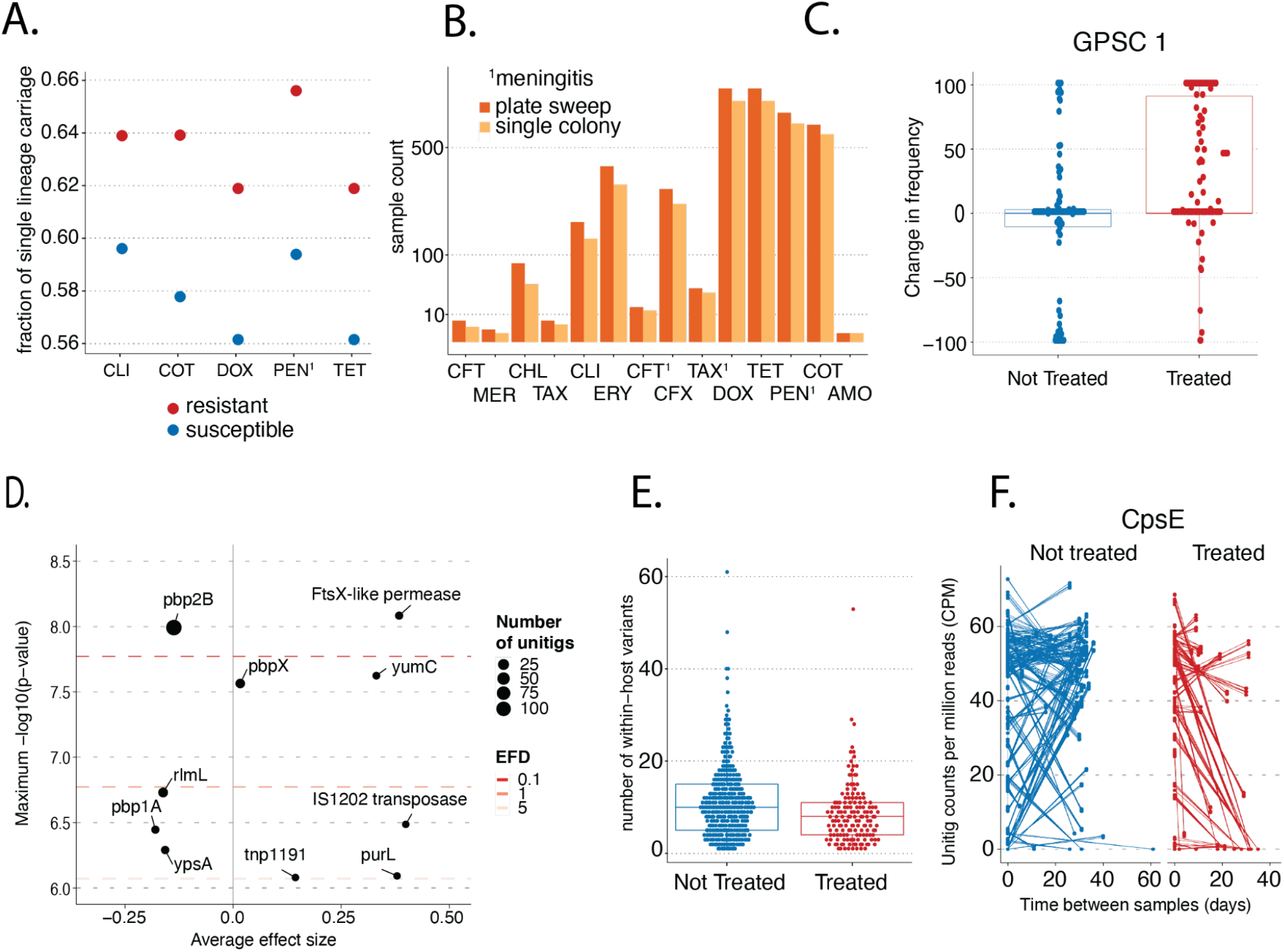
a.) The fraction of carriage events consisting of a single lineage found to be resistant to each antibiotic class. Only those classes found to be less likely to occur in instances of multiple colonisation than expected given the background prevalence in the population are shown. b.) The number of resistance calls for each antibiotic class in 1158 samples for which both single colony picks and PDS had been performed. c.) The distribution of the change in frequency of the GPSC1 lineage in pairs of consecutive samples that have and have not received antimicrobial treatment. d.) A dot plot indicating the significance and effect size of unitigs found to be associated with antimicrobial treatment. e.) The number of within-host SNV in samples involving only a single pneumococcal lineage split by those that have recently received antimicrobial treatment. f.) the normalised count of unitigs found in CpsE in pairs of samples where a subset had received treatment in between sampling events.

### Antibiotic treatment has a strong impact on pneumococcal within-host genetic diversity

We next considered selection in response to antimicrobial treatment, both in terms of the displacement of pre-treatment strains, and the microevolution of surviving pneumococci. Pairs of consecutive samples taken from the same infants within 100 days were selected where a subset had received antibiotics in between the sampling timepoints. The prescription of antimicrobials in the study participants was monitored by the study team and care was taken to document both antimicrobials prescribed by the SMRU clinic and those obtained from other sources (64).

The multidrug-resistant (MDR) lineage GPSC1 was found at considerably higher frequencies than other GPSCs following treatment (Figure 4C, Supplementary Figure 18). A similar analysis using the Penicillin Binding Protein (PBP) gene ‘types’ used in the *in-silico* classification of pneumococcal resistance by the CDC (65) identified the PBP2X-47 and PBP1A-13 types as being strongly associated with the persistence of a lineage post treatment (Supplementary Figure 19). These are the most common types in GPSC1 in Maela, representing 54% and 98% of the single colony isolates respectively (24). Alterations in the PBPs reduce their affinity for penicillin and thus susceptibility to beta lactam antibiotics while allowing them to maintain their role in cell wall metabolism.

Although it has been suggested that the increase in resistant isolates following treatment is due to the elimination of susceptible lineages (66), we were able to show that this pattern is still observed after controlling for presence of GPSC1 in the first sample of a pair. This suggests that not only does treatment increase the frequency of GPSC1 by eliminating competing lineages, it also increases the chance that a child will be colonised with the resistant lineage following treatment. Hence resistant bacteria also benefit from reduced displacement by competing strains after antibiotic consumption. This could motivate pre-emptive interventions, such as limiting contacts with high-risk individuals following antibiotic treatment.

To investigate selection acting within a single carriage event, we considered the set of paired samples where the same lineage was present in both samples of each pair. To incorporate variation in the accessory genome we looked at the frequency of variable length sequence elements (unitigs), as is common in bacterial GWAS studies (67, 68). This revealed that the overall diversity of within-host variants reduced markedly following antimicrobial treatment (Figure 4E). We used a linear model to test for the change in prevalence of a variant after antimicrobial treatment (Methods). This identified variants that were consistently found at lower frequencies post treatment including in the capsular gene cpsE (Figure 4F). Point mutations in *cpsE* have been shown to alter the growth, adherence and competence of pneumococci (69, 70). More common variants with smaller effect sizes were observed in the adenylosuccinate synthetase gene (*purA*), and genes involved in the zinc (*adcC*) and magnesium transport (*corA*) systems. The expression of all three of these genes has previously been observed to be downregulated in response to sub-inhibitory concentrations of penicillin (71). An explanation for this is that antimicrobial treatment induces a significant bottleneck in the pneumococcal population within the host even when the resident strain is resistant. This results in lower frequency or less fit genotypes being outcompeted and may be an example of short sighted bacterial evolution. Here, mutations that are beneficial within the host may have a reduced chance of being transmitted possibly due the increased chance of treatment induced clearance (72).

While the paired sample deep population GWAS can identify changes occurring within a single carriage event, it is not able to identify variation associated with selection against whole lineages. By comparing the presence and absence of unitigs in samples taken within 28 days of treatment to those that had not been treated and controlling for population structure using a Linear Mixed Model (68) we identified a number of sequence elements associated with antimicrobial treatment (Figure 4D). In keeping with the previous analysis, this included elements found in *pbp2B*, *pbp2X* and *pbp1A* — three of the genes that encode for the major penicillin binding proteins which are critical in determining non-susceptibility to beta-lactam antibiotics (73, 74). Interestingly, the strongest association was with *pbp2B,* which is the primary gene for low-level penicillin resistance, and is consistent with amoxicillin being used for treatment in (66.9%) of cases (75). The stronger association with *pbp2B* indicates that resistance conferred by these mutations are found across a diverse set of lineages, while the associations observed in the paired analysis of PBP types is driven primarily by particular lineages such as GPSC1. However, strong epistatic links have been observed between these genes as alterations in all three is necessary for pneumococci to develop broad range nonsusceptibility to beta lactam antibiotics (76).

We also observed associations with the membrane protein FtsX, a ribosomal RNA methyltransferase (*rlmL*) and a ligand binding protein (YpsA). FtsX is involved in cell division and is thought to co-locate with both PBP2b and PBP2x in the outer-ring peripheral peptidoglycan synthesis machine during cell division (77, 78). YpsA is also linked to pneumococcal cell division (79). RmlL is thought to facilitate resistance occurring through other mutations (80). A plausible explanation for these associations is that they allow pneumococci to slow down their metabolism and cell division, increasing the chances that the population is able to persist over the time period when the antibiotic is present.

Interestingly we also observed a weak association with the insertion sequence IS1202, which has been closely linked to the MDR associated serotype 19F and its capsular polysaccharide synthesis (*cps*) locus, which are predominantly found in GPSC1 (81). Critically, unlike the vast majority of studies which rely on laboratory resistance profiles to investigate the mechanisms that confer resistance, these associations were found directly within natural human carriage. This highlights the significant added contribution that deep population GWAS techniques can have in identifying pathogen mutations that may be associated with treatment failure.

## Discussion

Our ability to understand the within-host evolution and transmission of *S. pneumoniae* within the host is essential to developing successful public health interventions. We have shown that deep within-host population sequencing can lead to substantial improvements in surveillance of high-risk genotypes, reconstruction of transmission chains, and the impact of antibiotic resistance on co-colonisation and competition. In particular, we were able to double our sensitivity for detecting the highly-invasive serotype 1 in carriage. These lineages were often found at low frequencies which may explain the disconnect between their high prevalence in invasive disease and scarcity in carriage studies that rely on either latex sweeps or representative genomes. The increased resolution of PDS also revealed an age dependent rate of transmission between mothers and infants. This coupled with the strong association between geographic distance and the likelihood of direct transmission within the Maela refugee camp suggests that interventions targeting close contacts could be particularly important for reducing disease and colonisation by resistant lineages in early childhood prior to vaccination and following antimicrobial treatment.

Our results provide a large-scale dataset on the natural co-colonisation of both resistant and susceptible pneumococcal lineages within the same host. We provide clear evidence that such coexistence is frequent (previously an assumption made by a number of models (5, 6, 63)) and find that resistant lineages appear less often in multiple colonisation than expected given their overall frequency within the population, consistent with the lower fitness of resistant lineages observed in laboratory experiments. The negative association between resistance and multiple colonisation, combined with the association between antimicrobial treatment and subsequent colonisation by a multidrug resistant strain, indicates that reduced within-host competition following treatment plays a major role in the risk that an infant is colonised with an MDR lineage. This emphasises that the broader dynamics of pathogen population structure and interstrain competition must be a key consideration in the design of vaccines and other interventions (36). The observed competition could also motivate the use of preemptive probiotics to protect against colonisation by more dangerous lineages, although trials of such approaches have returned mixed results (82, 83). The strong negative selection observed in heat shock proteins suggests that multiple-antigen vaccines may provide a valuable alternative to current capsule specific vaccines as they have the potential to elicit cross-serotype protection (60). Overall, the added insights into selection and evolution within the host, coupled with the significant improvements in transmission inference and surveillance, presents a compelling case for the future routine use of deep within-host population sequencing in the research and surveillance of common bacterial pathogens.

## Methods

### Sample selection

Nasopharyngeal swabs were collected between November 2007 and November 2010 from an initial cohort of 999 pregnant women leading to the enrolment of 965 infants. From 23910 swabs collected during the original cohort study, 19359 swabs from 737 infants and 952 mothers were processed according to World Health Organisation (WHO) pneumococcal carriage detection protocols (84) and/or the latex sweep method (85). All isolates were serotyped using latex agglutination as has previously been described (14).

Deep sequencing of sweeps of colonies was attempted on a subset of 4000 swabs. All swabs taken before and after an antibiotic treatment event were selected. Further swabs were included if they were inferred to be within close transmission links corresponding to a distance of (<10 SNPs) using a previously sequenced set of 3,085 whole genome sequences obtained from single colony picks (24).This allowed for increased resolution into both the impact of antibiotic treatment on within host diversity and consideration of the transmission bottleneck. A subset of 25 mother/child pairs were also sequenced at a higher temporal resolution of at least once every 2 months. The remaining samples were selected randomly.

### Culture and sequencing

Specimens, 100 μl of nasopharyngeal swab stored at −80°C in skim milk, tryptone, glucose and glycerin media, were plated onto Columbia CNA agar containing 5% sheep blood (BioMerieux, product reference 43071). These were incubated overnight at 37±2 °C with 5% CO_2_. All growth was collected using sterile plastic loops and placed directly into Wizard Genomic DNA purification kit Nuclei Lysis solution (Promega, reference A1120). The Wizard kit extraction protocol was then followed, eluting in 100 μl of the provided DNA Rehydration Solution. DNA was quantified with a BioPhotometer D30 (Eppendor) and then stored at −80°C prior to sequencing. DNA extractions were sequenced if they contained more than 2.5 μg of DNA. Sequencing was performed at the Wellcome Sanger Institute on an Illumina NovaSeq at 192 plex using unique dual index (UDI) tag sets.

### Quality control filtering

In total, 3961 samples were successfully sequenced, including 200 which were sequenced in replicate. In order to concentrate our efforts on those samples where there was sufficient data to retrieve reliable results we excluded samples with a mean coverage below 50 fold representing 20% of the median coverage observed across all samples (Supplementary Figure 8). Whilst it is hard to choose an optimal coverage threshold, 50x has been shown to be a reasonable coverage for the assembly of bacterial genomes (86). To account for contamination from other species, Kraken was run on all samples with a histogram of the proportion of each sample assigned to *S. pneumoniae* given in Supplementary Figure 9. A threshold of requiring that at least 75% of reads were classified as *S. pneumoniae* was chosen as a compromise between avoiding excluding too many samples and ensuring contamination did not bias our analyses. Further checks were also conducted at each stage of the downstream analyses to ensure results were not impacted by remaining low levels of contaminating species. Overall, this resulted in 3188 samples including 164 replicates that were considered in the subsequent analysis steps.

### Lineage deconvolution

Lineage deconvolution was performed via the mSWEEP and mGEMS algorithms (87, 88) using a reference database consisting of a high quality subset of 20,047 genomes from the Global Pneumococcal Sequencing Project database (9). Included in this subset were 2,663 genome assemblies from the original genome sequencing study of the Maela camp that relied on single colony picks (24). The PopPUNK algorithm was used to assign each of these genomes to their respective Global Pneumococcal Sequencing Cluster (89). The mSWEEP and mGEMS pipelines were then run using the fastq files for each deep sequencing sample with the exact commands used given in the Rmarkdown provided as part of the accompanying GitHub repository (87, 88). To reduce the possibility of false positives lineages were only called if they were present at a frequency of at least 1%. The Mash Screen algorithm was also run on each of the deconvoluted lineages using the same reference database (90). Only lineages that shared at least 990/1000 hashes were retained.

### Serotype calling

Serotypes were identified by taking the union of two pipelines (Supplementary Figure 2). The serocall algorithm was run on the raw fastq files for each sample (25). As a second step, the seroBA algorithm was run on each of the deconvoluted lineages identified by mGEMS pipeline (91). By comparing the results of these pipelines on both artificial laboratory mixtures (25) and samples for which single colony picks had also been performed we were able to determine that while both algorithms generally agreed at the serogroup level the serocall algorithm was more sensitive and was able to detect lineages below the 1% cut-off used in running mGEMS. As the serocall algorithm was less precise at distinguishing serotypes at the sub-group level (Supplementary Figure 11), whenever the pipelines produced conflicting results at the sub-serogroup level the seroBA result was chosen. After taking the union of these two pipelines we were able to correctly recover 93.6% of serotypes originally identified by latex sweeps performed on the same set of samples. The analysis of the artificial laboratory mixtures also indicated that the combined pipeline achieved a sensitivity of 0.93 with a precision of 1.

### Resistance calling

Similar to the calling of serotypes, resistance determinants were identified via two pipelines using the raw data and the deconvoluted output of the mGEMS pipeline. The pneumococcal specific CDC resistance calling pipeline was run on each of the deconvoluted lineages identified using mGEMS (65, 92). This makes use of a database of PBP proteins with known resistance profiles. The combined mGEMS and resistance calling pipeline was found to achieve a sensitivity of 0.75 and precision of 0.825 in identifying resistance calls from the artificial laboratory mixtures. The lower accuracy in identifying resistance was caused by small inaccuracies in the deconvolution of strains and a lower sensitivity in detecting resistance in the sample containing 10 lineages. As the maximum number of lineages observed in any sample in our dataset was six this drop in sensitivity at very high multiplicities of infection did not present a significant problem. To account for inaccuracies in the deconvolution of resistance associated sequencing reads we only report resistance calls at the sample level. After restricting the comparison of laboratory calls to those samples continuing <10 lineages we achieved an accuracy of 1 at the sample level. To verify the pipeline on a more diverse dataset we compared the resistance calls found in 1158 samples for which both single colony picks and whole plate sweeps had been taken. The mGEMS + CDC pipeline was able to achieve a recall rate of 96.9% indicating that the combined pipeline is able to accurately identify resistance from deep sequenced plate sweeps. To check that the pipeline did not result in a high number of false positives we compared the calls from single colony picks and plate sweeps on the subset of 584 samples which involve only a single lineage. Here, we would expect the results of both approaches to be similar. Supplementary Figure 16, indicates that there was not a significant difference on this subset of samples with only a very small increase of 2.7% (53/1980) of resistance calls (p=0.4, poisson GLM).

### Resistance co-occurrence

To examine whether certain lineages or serotypes were more likely to be found in instances of multiple colonisation we performed a logistic regression using a generalised linear mixed model with a complementary log-log link function. To control for the increase in the probability of resistance being present in a sample with multiple lineages simply because there were more lineages present we used an offset term. This is a common approach used in ecological studies to control for the differences in exposure when investigating a binary outcome. This allows us to test whether the presence of resistance as a binary dependent variable is associated with multiple colonisation beyond what would be expected given the background frequency of resistance in the population.

To control for the lineages present within each sample we performed Multidimensional Scaling (MDS) on a pairwise distance matrix inferred using the Mash algorithm between each sample (93). The first ten components were included in the regression to control for population structure as is common in bacterial GWAS studies (94). Host effects were controlled for by including a random effect for the host.

### Genome wide association analyses

To better account for the extensive pangenome in *S. pneumoniae*, locus level association analyses were performed using an alignment free method which first identifies all unique unitigs (variable length k-mers) within the samples being considered. Unitigs have been shown to better account for the diverse pangenomes found in bacteria (67). The frequency of each unitig in each sample was obtained by first running the Bifrost algorithm to define the complete set of unitigs present (95). The count of each unitig in each sample was then obtained using a custom python script available in the accompanying GitHub repository. To avoid testing very rare features, we only considered those unitigs present in at least 1% of the samples of interest in our presence/absence based analysis and in at least 2% of our paired analysis discussed below.

To investigate the impact of antibiotic treatment on *S. pneumoniae* carriage we performed two main analyses. The first consisted of a typical case control design and compared samples that were within a recent antimicrobial treatment event to those where no treatment had occurred. This allowed us to investigate features associated with recent antibiotic treatment but does not consider the changes that occur within an individual that is already colonised with *S. pneumoniae* prior to treatment. In order to shed light on this scenario our second analysis investigated the impact of treatment on pairs of consecutive samples from the same patient where a subset of patients had received antibiotic treatment in between samples (Supplementary Figure 12).

#### Standard design

Samples were classified as treated if they were within 28 days of an antimicrobial treatment event. This was chosen after reviewing the decline in the proportion of resistant isolates tested via disk diffusion and Etest MIC testing of all swabs positive for *S. pneumoniae* (Supplementary Figure 13). The python implementation of the seer algorithm was then used to identify unitigs significantly associated with treatment (68). Here, rather than using counts unitigs were called as either present or absent. To control for population structure Pyseer was run using a Linear Mixed Model with a kinship matrix generated by taking the cross product of the binary unitig presence/absence matrix. Unitigs found to be significant were then aligned to a collection of pneumococcal reference genomes including all the single genome assemblies of Chewapreecha et al., (24) and assigned a gene annotation based upon the reference gene in which they aligned. Only those unitigs that were successfully aligned were considered for further analysis. To account for the large number tests performed we considered three p-value thresholds corresponding to an Expected number of False Discoveries (EFD) of 0.1, 1 and 5. The 0.1 threshold corresponds with the commonly used Bonferroni method while the more relaxed thresholds allowed us to consider weaker signals. All three of these thresholds were more stringent than controlling for the False Discovery Rate using q-values which has been suggested as an alternative to the Bonferroni method as it is often found to be overly conservative (96). Combined with past knowledge of likely resistance elements in *S. pneumoniae* we were able to confidently identify associations.

#### Paired design

Our unique sampling allows us to compare samples from the same individual before and after treatment. We first identified sample pairs where there were at most 100 days separating pneumococcal positive nasopharyngeal swabs from the same individual. We restricted our analysis to infants as treatment information for mothers was not available. To ensure previous treatments prior to the first sample of an individual were not confounding our results we excluded pairs with any treatment event within 28 days of the first swab. This resulted in 615 sets of paired samples. We classified these pairs into treated and untreated groups based on whether or not the individual had received antibiotic treatment in the time between swabs.

We only considered paired samples where the infants were positive for *S. pneumoniae* in both samples. As a result we are not considering the impact of antibiotic treatment on overall carriage rates but rather the differences in *S. pneumoniae* genomes pre and post antibiotic treatment. Using this paired design we considered the impact of treatment both at the lineage (GPSC) level as well as the locus level. This distinction between lineage and locus is similar to that defined in (97). Unlike many previous bacterial GWAS studies which typically focus on the presence or absence of a feature, we considered the frequency of both lineages and loci within each sample. This improves our ability to identify more subtle changes that can be obscured by ignoring within-host diversity.

#### Lineage level

At the lineage level we considered the estimated frequencies of each lineage obtained using the mSWEEP algorithm. We used a simple linear model to test whether treatment impacted the frequency of the second sample of a pair after controlling for the observed frequency in the first sample as well as the difference in time between the two samples.

#### Locus model

To investigate locus level effects we considered the frequency of each unitig in each sample. In order to control for lineage level effects we concentrated on pairs where the same lineage was present in both samples. This reduced the analysis to 445 pairs.

Unlike the lineage level analysis where we used estimated frequencies, unitigs are represented by the number of times they were observed in the raw reads from each sample. This is a similar problem to that found in the analysis of RNAseq datasets where the number of RNA reads aligned to a gene is used as a proxy for the expression of that gene. Using a similar approach to that commonly used in the analysis of RNAseq data we fit a linear model to the log unitig counts normalised by the number of reads sequenced in each sample. Similar to the commonly used ANCOVA method for analysing pre and post treatment data we used the pre-treatment count to control for the paired nature of the data (98). We also include a covariate to control for the time between when the samples were taken. Further explanation and the code used to run all the association analyses is available in Supplementary Text included in the GitHub repository.

### Within-host variant calling

To identify within-host variants we ran the LoFreq variant calling pipeline on all samples for which only a single GPSC lineage had been identified with mSWEEP. The Lofreq pipeline has been shown to generate robust minority variant calls and accounts for base call qualities and alignment uncertainty to reduce the impact of sequencing errors (99, 100). To mitigate the impact of reference bias, each sample was aligned to a representative assembly (the medoid) for the GPSC that it most closely resembled via Mash distance (93). Reads were aligned to the chosen reference genomes using BWA v0.7.17-r1188 (101). The Picard tools v2.23.8 ‘CleanSam’ function was then used to soft clip reads aligned to the end of contigs and to set the alignment qualities of unaligned reads to zero. Pysamstats v1.1.2 was run to provide allele counts for each location of the aligned reference for use in the transmission analysis (102). The LoFreq pipeline v2.1.5 was run initially with stricter filters requiring a coverage of at least 10 reads to identify a variant. The resulting variant calls were used along with the read alignment as input to the GATK BaseRecalibrator tool v4.1.9 as suggested in the LoFreq manual to improve the estimated base quality scores (103). Finally the LoFreq pipeline was run for a second time with a reduced coverage requirement of 3 reads. The resulting variant calls were only considered if there was support for the variant on at least two reads in both the positive and minus strand. Of the remaining within-host single nucleotide variants, Supplementary Figure 4 indicates that there was strong agreement between variant calls in the set of 95 sequencing replicates for which only a single lineage was present with a median of 91.7% of variants recovered. The distribution of minority variants among different coding positions was also consistent with real mutations rather than sequencing errors with variants at the third codon position being most frequent (Supplementary Figure 5) (104).

### Filtering problematic regions

To identify problematic variants that were likely the result of low level contamination or multi-copy gene families we implemented a similar approach to that used to identify recombination in the tool Gubbins (105). A scan statistic was used to identify regions of the alignment with an elevated number of polymorphisms. Assuming that within host variants are relatively rare and should be distributed fairly evenly across the genome, regions with a high number of polymorphisms are likely to be the result of confounding factors and can thus be filtered out.

We assumed a null hypothesis *H*_0_, that the number of polymorphisms occurring in a window *s*_*w*_ follows a binomial distribution based upon the number of bases within the window *w* and the mean density of polymorphisms across the whole alignment. We chose *w* for each sample such that *E*(*s*_*w*_) = 1. A window centred at each polymorphism was then considered and a one-tailed Binomial test was performed to determine if that window contained an elevated number of polymorphisms. After adjusting for multiple testing using the Benjamini-Hochberg method those windows with a p-value < 0.05 were selected and combined if they overlapped with another window (106).

To more accurately define the edges of each region, we assumed each combined window conformed to an alternative hypothesis *H*_1*r*_ where the number of polymorphisms *s*_*r*_ also followed a binomial distribution with a rate based on the length of the window *l*_*r*_ and the number of polymorphisms within the window *s*_*r*_. Each end of the window is then progressively moved inward to the location of the next polymorphism until the likelihood of *H*_1*r*_ relative to *H*_0_ no longer increases. The resulting final windows were then called as potential problematic regions if they satisfied the inequality

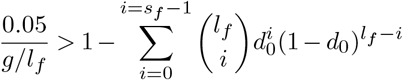

Where *l*_*f*_ is the length of the final w indow, *g* i s t he length of the reference genome and *d*_0_ is the expected rate of polymorphisms under the null hypothesis. The left hand side of the equation accounts for the possible number of similarly-sized non-overlapping windows in the reference. To further reduce the chance that spurious alignments between homologous genes could bias our results we took a conservative approach and excluded mutations which were found within a single read length (150bp).

### Mutational spectrum

In the mutational spectrum analysis of human cancers, normal samples are usually taken along with samples of the cancer to allow for somatic mutations to be distinguished from germline mutations. As we can not be sure which alleles were present at the start of a pneumococcal carriage episode we can not be certain of the direction a mutation occurred in. For example, it is difficult to distinguish between an A−>C and a C−>A mutation. Instead, we consider the difference between the consensus and minority variants at each site in the reference genome. If we assume that the colonising variant typically dominates the diversity within an infection then this approach corresponds with the direction of mutation. To account for the context of each mutation we considered the consensus nucleotide bases on either side of the mutation. These were then normalised to account for the overall composition of the reference genome for each GPSC. The normalised mutation rates (*r*) for each of the 192 possible changes (*j*) in a trinucleotide context were calculated as:

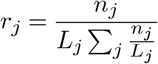

Where *n*_*j*_ is the total number of mutations observed for a trinucleotide change *j*, and *L*_*j*_ is the total number of times that the corresponding trinucleotide is present in the reference genome. To avoid double counting the same mutation, each variant was only counted once per host. The resulting frequencies for within and between hosts are given in Supplementary Figure 17. The frequencies of each of the single nucleotide changes without accounting for sequence context were calculated similarly.

To compare with the mutational spectrum observed across a longer timescale we considered the recombination filtered alignments of 7 major sequence clusters generated in the original publication of the single colony pick analysis of the Maela dataset (24). We used Iqtree v2.1.2 to build a maximum likelihood phylogeny for each alignment using a General Time Reversible (GTR) model with 4 rate categories and enabled the ‘ancestral’ option to reconstruct the sequences at the internal nodes of the resulting phylogeny (107). Mutations were called by considering changes in alleles between consecutive nodes of the phylogeny and the mutational spectrum was normalised using the trinucleotide frequencies in the reconstructed ancestral sequence of the root node. A permutation test was used to compare the propotion of each mutation type found in the within-host and between-host sets.

### Selection

Selection analyses were performed using a modified version of the dNdScv package (53) to allow for the incorporation of variants called against multiple reference genomes. Distinct from traditional approaches to estimating dN/dS ratios which were developed to investigate selection in diverse sequences and rely on Markov-chain codon substitution models (108), dNdScv was developed to compare closely related genomes such as those found in somatic mutation studies where observed changes often represent individual mutation events. To avoid false signals of negative or positive selection which have been observed under simpler models (53), dNdScv uses a Poisson framework to account for the context dependence of mutations, non-equilibrium sequence composition and to provide separate estimates of dN/dS ratios for missense and nonsense mutations.

To extend dNdScv to allow for the use of multiple reference genomes we first clustered the gene regions from the annotated reference genomes using Panaroo v1.2 (109). The impact of each of the mutations identified using the LoFreq pipeline was inferred using dNdScv for each sample separately using the corresponding reference genome and gene annotation file. The combined calls for each orthologous cluster were then collated and the collated set used to infer genome wide and gene level dN/dS estimates using a modified version of dNdScv available via the GitHub repository that accompanies this manuscript. We used the default substitution model in dNdScv, which uses 192 rate parameters to model all possible mutation in both trends in a trinucleotide contact as well as two *ω* parameters to estimate the dN/dS ratios for missense and nonsense mutation separately. Due to the large number of samples we used the more conservative dNdSloc method which estimates the local mutation rate for a gene from the synonymous mutations observed exclusively within that gene (110). Care is needed when interpreting dN/dS ratios estimated from polymorphism data as they can be both time dependent, providing weaker signals of selection for more recent changes and can be biased by the impacts of recombination (111, 112). However, these are unlikely to cause significant issues in this analysis as the short time scales involved mean that recombination is unlikely to occur at a rate sufficient to bias the results and as each variant call is derived at the sample level rather than by the comparison of two separate samples as is typically the case in dN/dS studies relying on multiple sequence alignments of diverse sequences. As an extra precaution we also excluded gene clusters identified as paralogous by the Panaroo algorithm to reduce the chance that spurious alignments between paralogous genes could bias the results.

### Transmission inference

To identify the likelihood of transmission between each pair of hosts we extended the TransCluster algorithm to account for genetic diversity within the host and to be robust to deep sequencing data involving multiple lineages.

The TransCluster algorithm expands the commonly used approach of using a SNP distance threshold to exclude the possibility of direct transmission to account for both the date of sampling and the estimated epidemiological generation time of the pathogen (38). However, hypermutating sites, contamination, sequencing error, multi-copy gene families and multiple colonisation all present additional challenges when investigating transmission using within-host diversity information (19, 41).

To account for these challenges we took a conservative approach and estimated the minimum pairwise SNP distance that could separate any pair of genomes taken from two samples. Thus, two samples were only found to differ at a site if none of the alleles in either sample at that site were the same (Supplementary Figure 7). To allow for variation in sequencing depth across the genome we used an empirical Bayes approach to provide pseudocounts for each allele at each site informed by the allele frequency distribution observed across all sites. A Multinomial-Dirichlet distribution was fit to the allele counts for each sample independently via the maximum-likelihood fixed-point iteration method (113). The inferred parameters were then used as pseudocounts and a frequency cut-off corresponding to filtering out variants less than 2% was used. All variant calls that were observed were retained. An implementation of the TransCluster algorithm including the code to account for varying sequencing depth is available from (114).

The estimated minimum SNP distance was then used as input to the TransCluster algorithm assuming a mutation rate of 5.3 SNPs/genome/year and a generation time of 2 months. These values were inferred using an adapted version of the TransPhylo algorithm on the previously sequenced single colony picks from the Maela camp (see Supplementary Methods included in the accompanying GitHub repository) (24). The estimated substitution rate conforms with previous studies investigating short term evolutionary rates in *S. pneumoniae* (18) and the estimated generation time is consistent with previous estimates of pneumococcal carriage durations and a uniform distribution of transmission events (57). This resulted in estimates of the most likely number of intermediate hosts separating two sequenced pneumococcal samples.

These estimates were then combined with epidemiological and serological information to identify the most likely direction of transmission between mothers and their children as is described in the main text.

## Data and materials availability

Supplementary code and meta data is available from https://github.com/gtonkinhill/pneumo_withinhost_manuscript. To protect the anonymity of study participants some epidemiological data has been obscured in the publicly available files. The original metadata files are available on request via the MORU Tropical Health Network Data Access Committee https://www.tropmedres.ac/units/moru-bangkok/bioethics-engagement/data-sharing. The transmission clustering implementation is available at https://github.com/gtonkinhill/fasttranscluster. The modified version of the dndscv algorithm is available at https://github.com/gtonkinhill/dndscv. Raw sequencing data is stored with the ENA under accessions given in Supplementary Table 1.

## Funding

Wellcome [206194 to S.D.B, 216457/Z/19/Z to C.C]; Wellcome PhD Scholarship Grant [204016/Z/16/Z to G.T.H]; Norwegian Research Council FRIPRO [299941 to G.T.H]; ERC [742158 to J.C.]. The Shoklo Malaria Research Unit is part of the Wellcome Trust Mahidol University Oxford Tropical Medicine Research Unit, which is funded by the Wellcome Trust [220211];

## Acknowledgements

We thank all the laboratory staff at Shoklo Malaria Research Unit (SMRU) involved in performing culture and DNA extraction activities and staff at the Wellcome Sanger Institute that performed the sequencing. We thank Microsoft Research for assisting with resources used to run the deconvolution pipeline.

## Supplementary Figures

**Supplementary Figure 1.**
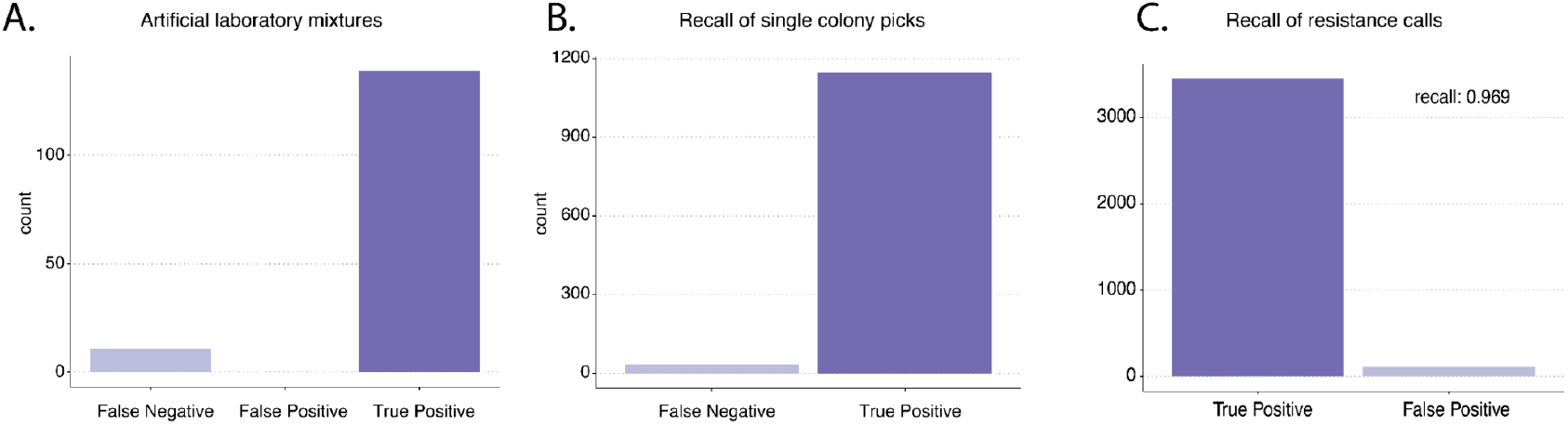
The number of true positive and false negative GPSC lineage calls in a.) 44 artificial laboratory mixtures from Knight et al., (25) and b.) 1158 samples which also had WGS performed on single colony pick in Chewapreecha et al., (24). c.) the recall of resistance calls in the same 1158 samples.

**Supplementary Figure 2.**
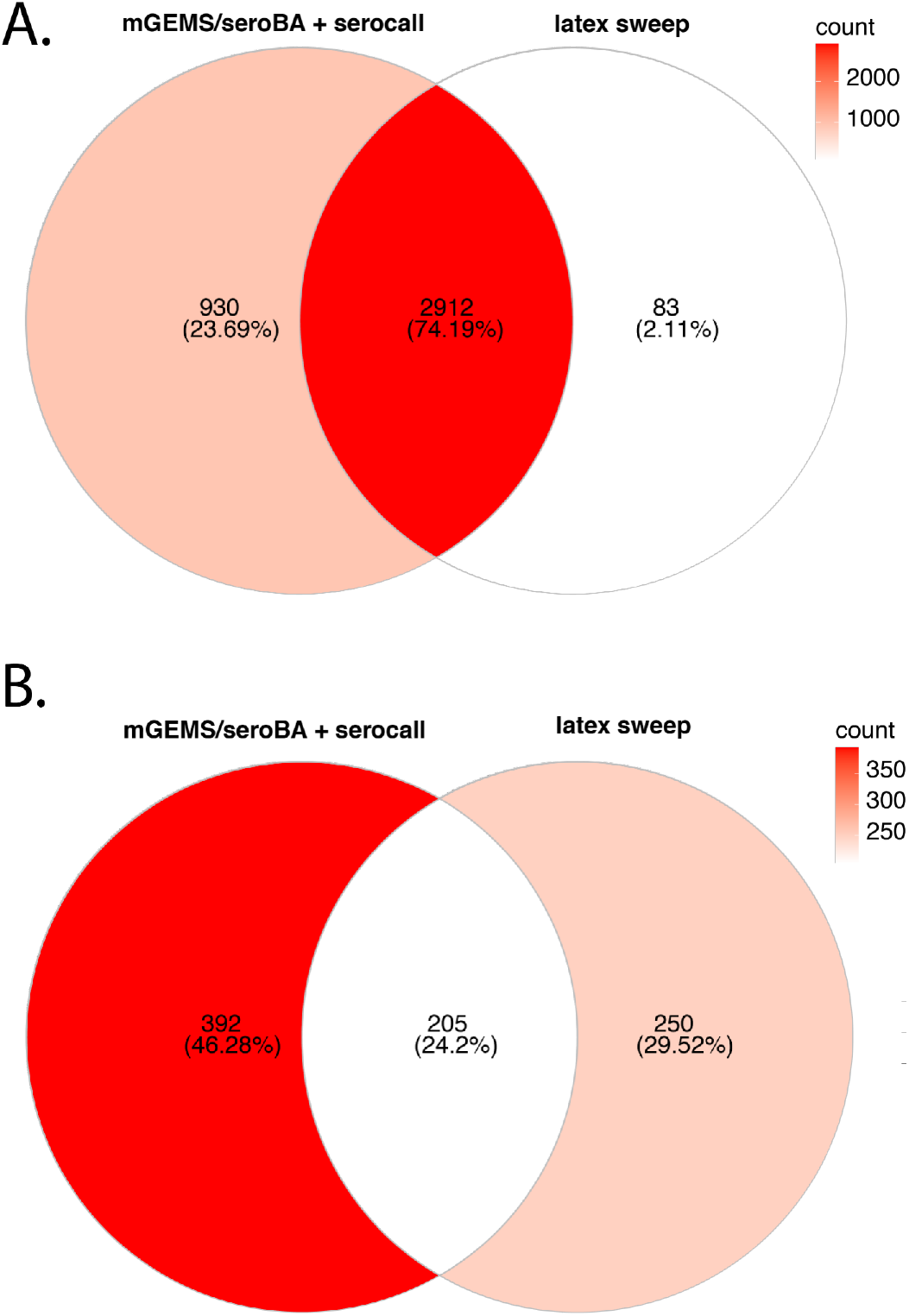
Intersection between the serotype calls of latex sweeps and the combined mGEMs/seroba and serocall pipeline for a.) all typable isolates and b.) non-typable.

**Supplementary Figure 3.**
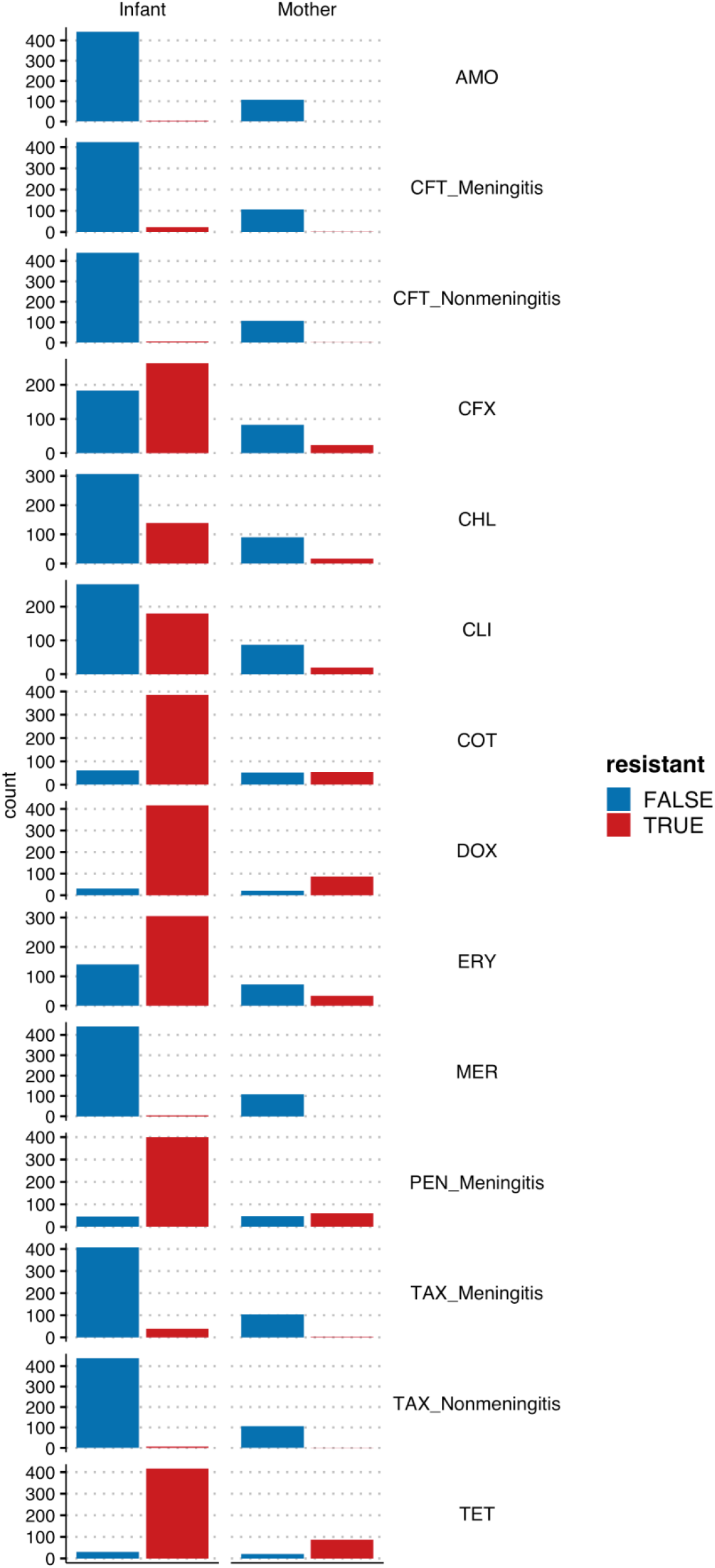
The number of samples found to be either resistant or susceptible to each antibiotic class for both mothers and infants. Resistance was determined by running the CDC pneumococcal resistance pipeline on the deconvoluted lineages output by the mGEMS pipeline. The individual lineage calls were collapsed to the sample level so that a sample was called as ‘resistant’ if resistance was observed in any of its lineage.

**Supplementary Figure 4.**
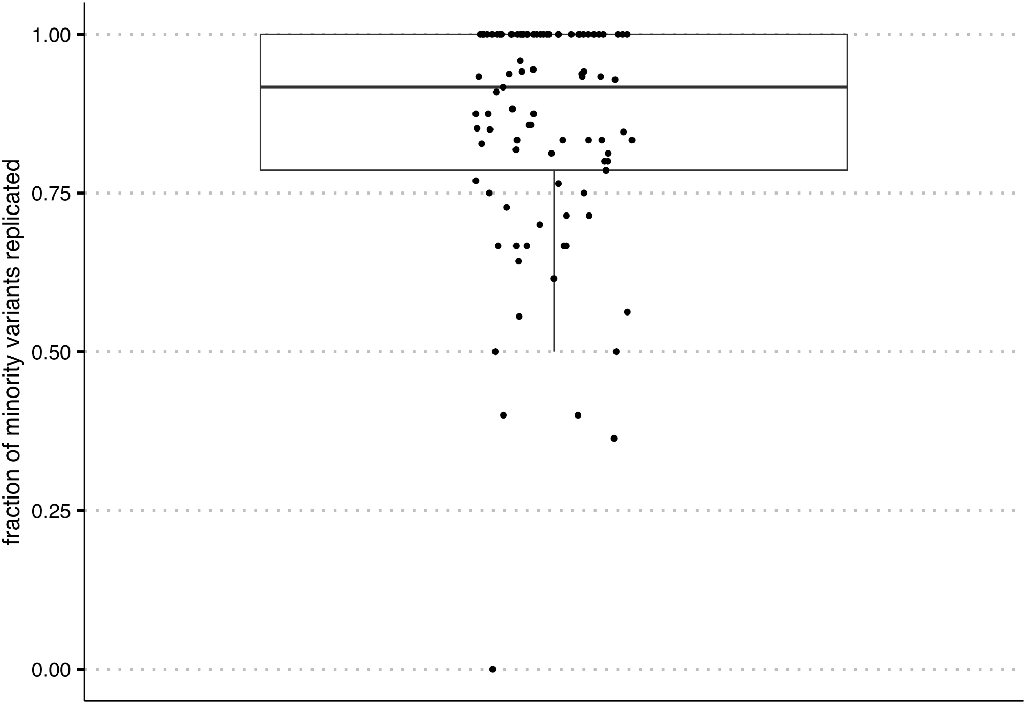
The fraction of minority single nucleotide variant calls replicated in 95 samples which involve only a single pneumococcal lineage and were sequenced in replicate with separate reverse transcription, PCR amplification, and library preparation steps.

**Supplementary Figure 5.**
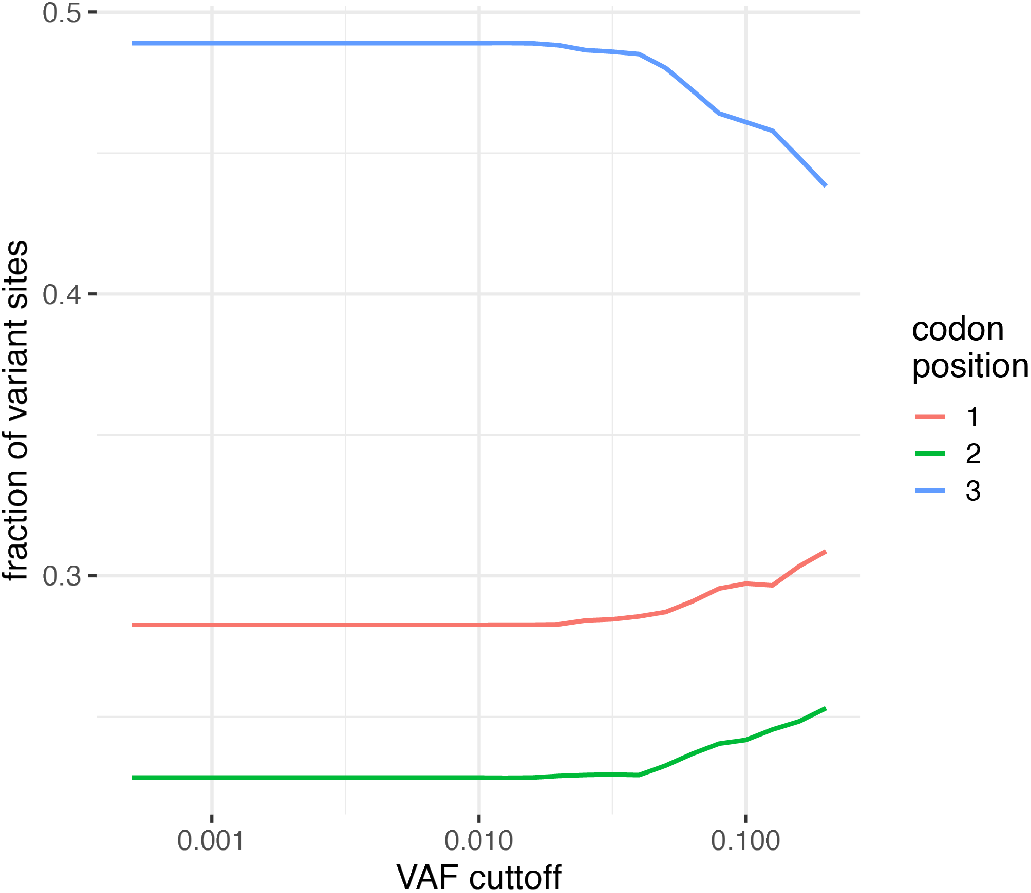
The distribution of the number of variable sites among different coding positions. Variable sites are dominated by those seen at the third codon position similar to that observed in Dyrdak et al., 2019. The stability of the fractions at lower frequencies suggests that the variant calling pipeline has successfully filtered out erroneous variant calls. At higher frequencies, the reduction in the total number of variants leads to increased variability.

**Supplementary Figure 6.**
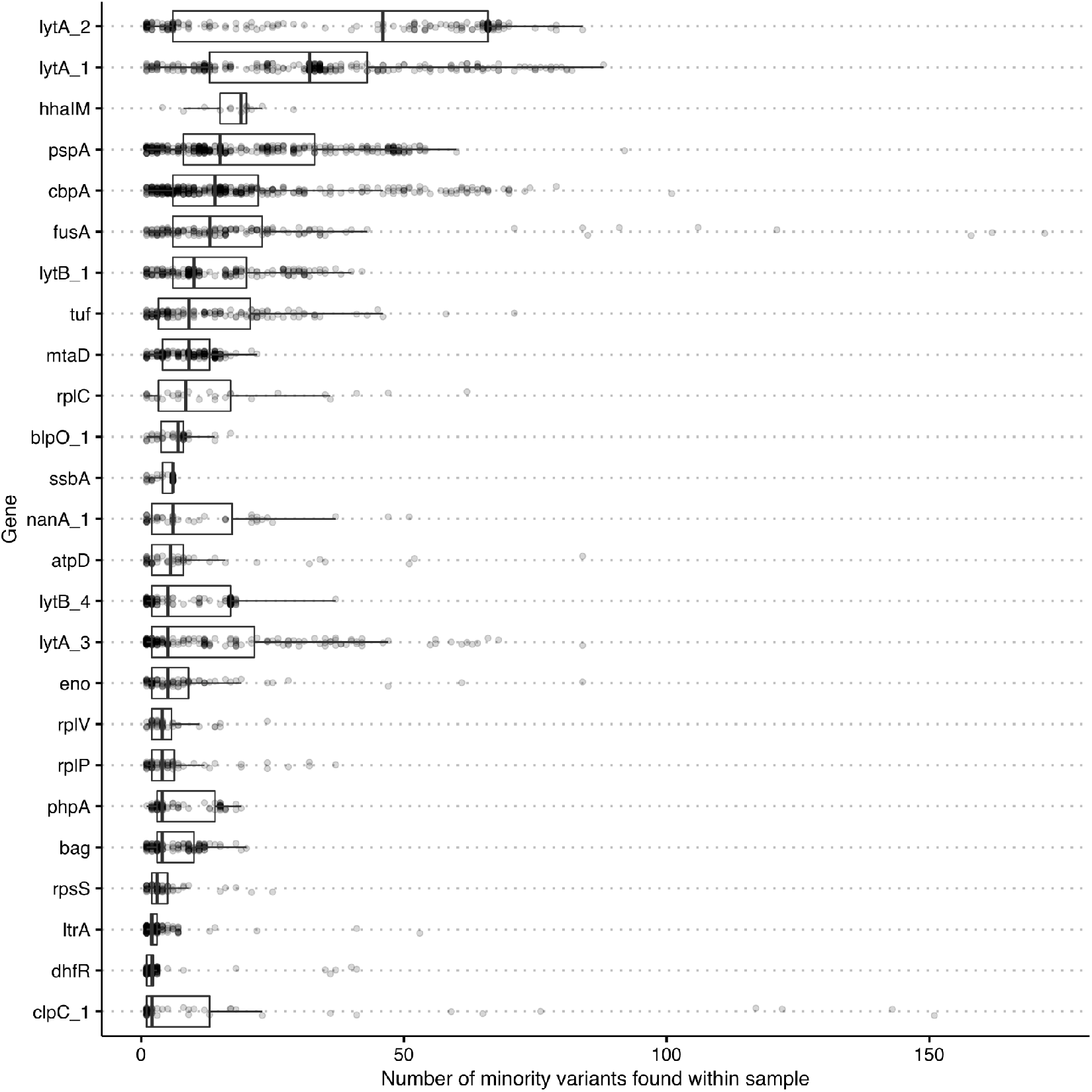
The distribution of the number of SNV found in regions with elevated rates of polymorphism classified by the spatial scan statistic (see Methods). The high rate of polymorphisms in these region indicates that these SNVs are unlikely to be the result of denovo mutation within the host and are instead likely to be driven by recombination, gene duplication, homology with phages and co-colonising bacterial species and hard to align regions.

**Supplementary Figure 7.**
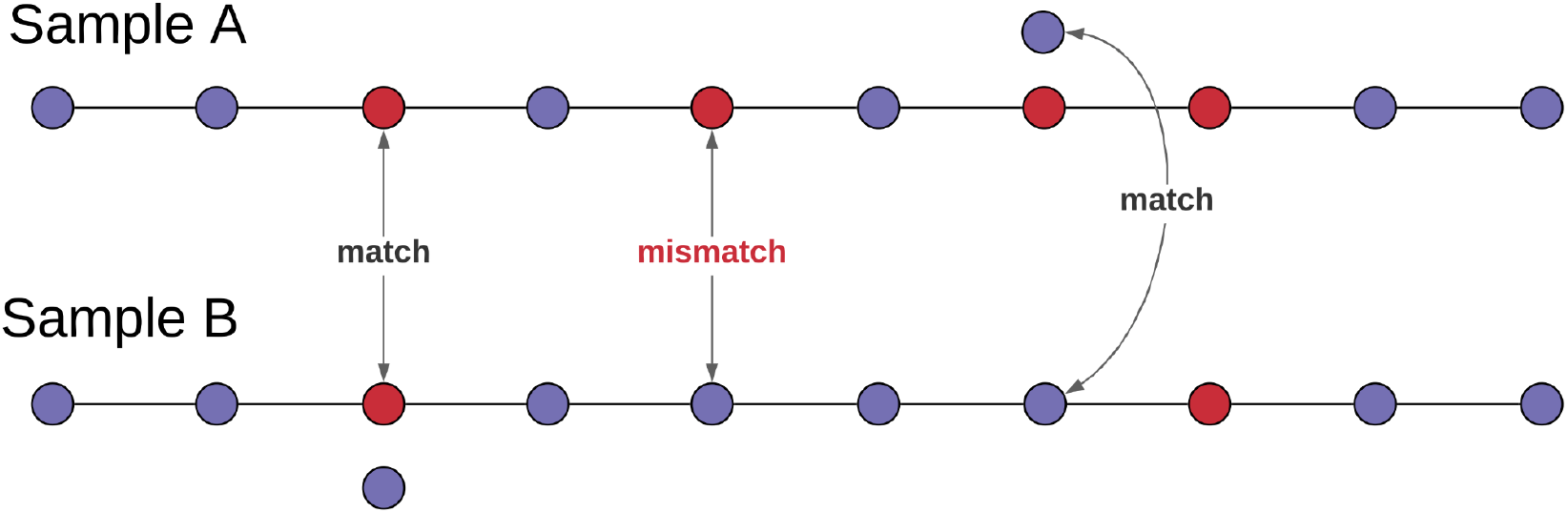
A schematic indicating how the pairwise SNP distance is calculated to account for within-host diversity and polymorphisms. Here, the red and blue indicate distinct nucleotides. A mismatch is only called if no alleles match at that location between the two samples. Variable sequencing coverage is accounted for using an empirical Bayes approach that made use of the multinomial Dirichlet distribution (methods).

**Supplementary Figure 8.**
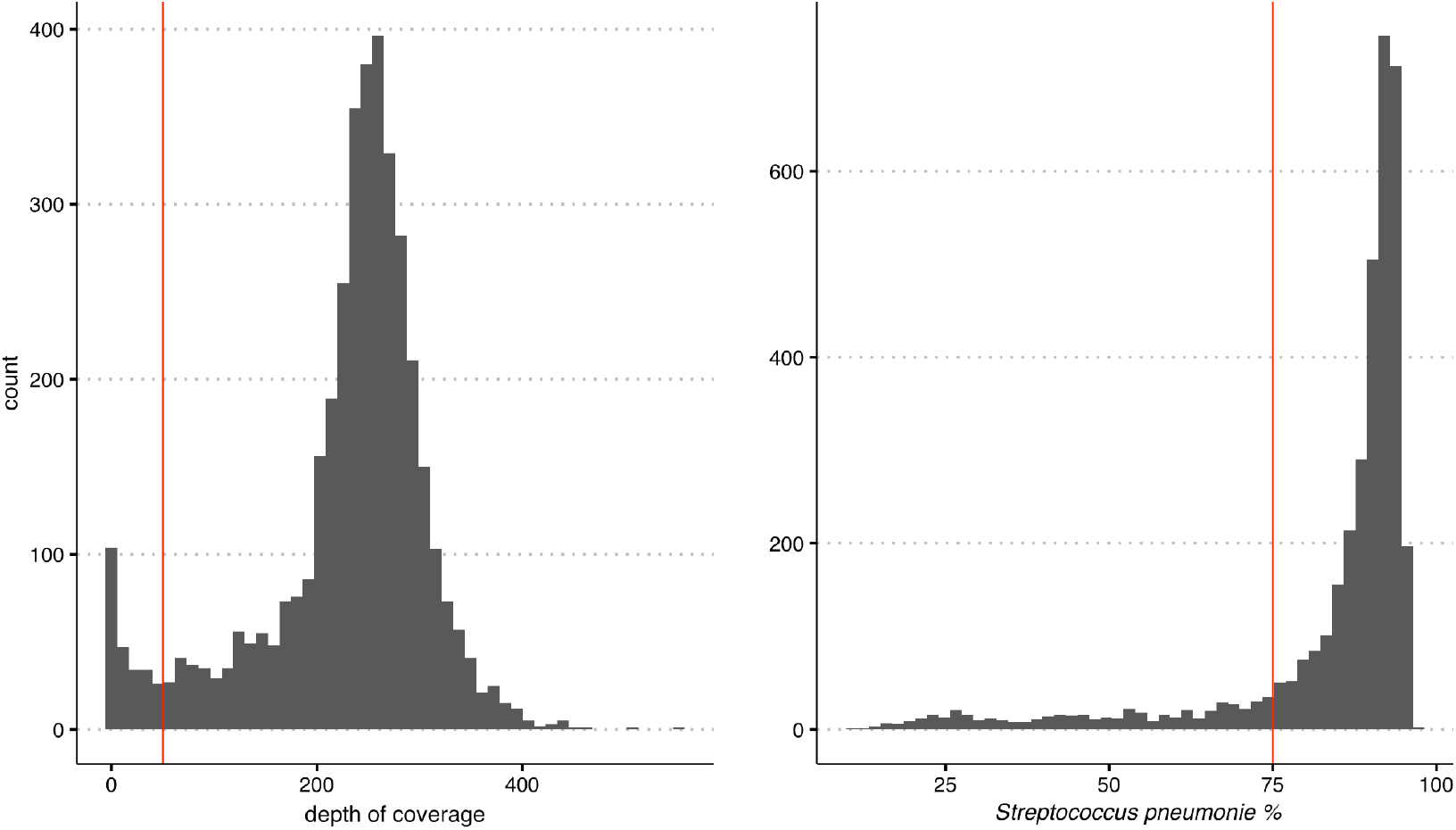
The distribution of the depth of sequencing coverage and fraction of reads that aligned to *S. pneumoniae* using the Kraken2 metagenomics read classification algorithm. The vertical red lines indicate the minimum thresholds chosen for samples to be included in the main analysis.

**Supplementary Figure 9.**
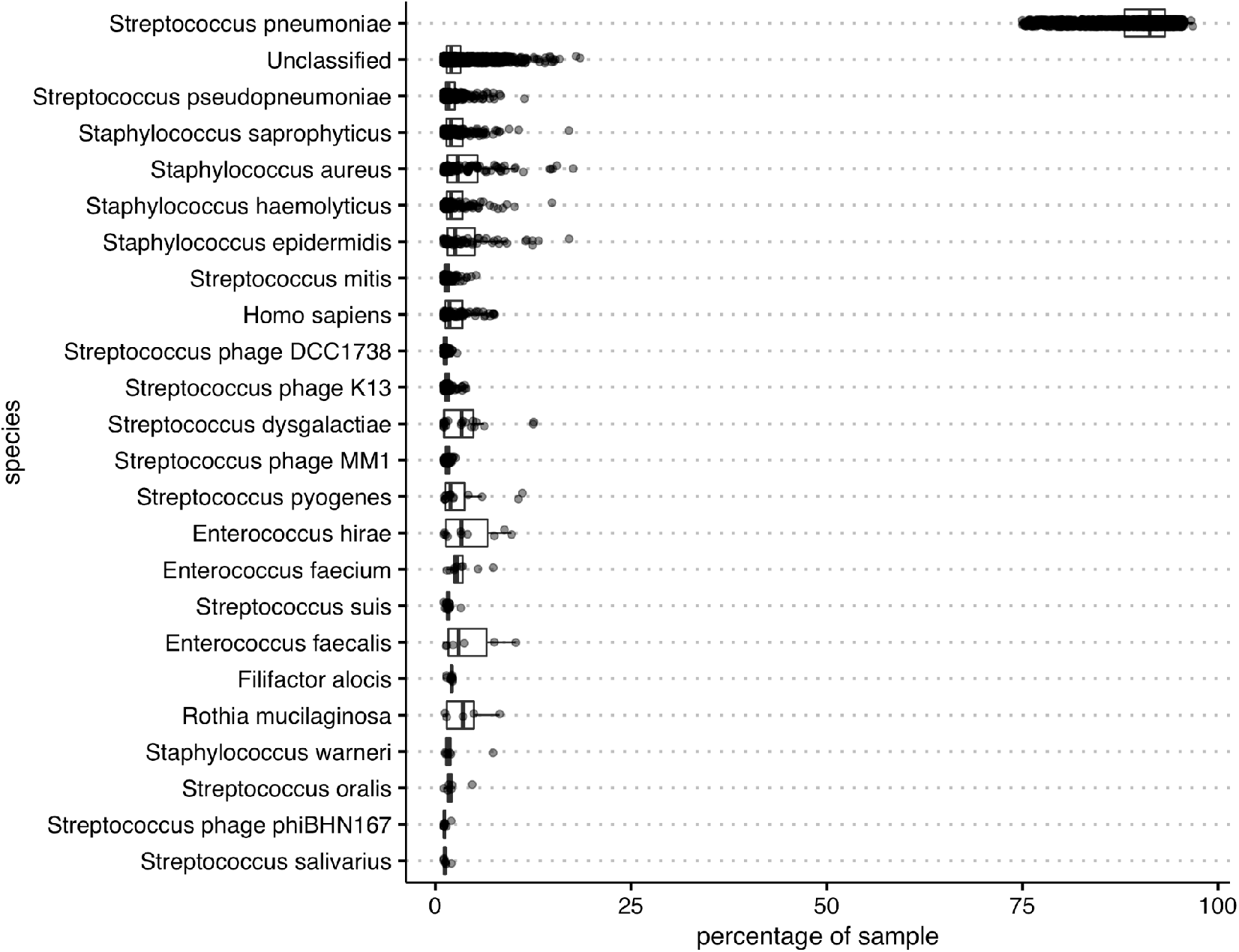
The distribution of the fraction of reads assigned to each species in each sample by the Kraken2 metagenomics read classification algorithm. Due to the large sequence diversity within, and similarity between, *S. pneumoniae* and *S. pseudopneumoniae*, a large fraction of reads assigned as ‘unclassified’ and as ‘*S. pseudopneumoniae*’ may actually belong to *S. pneumoniae* genomes.

**Supplementary Figure 10.**
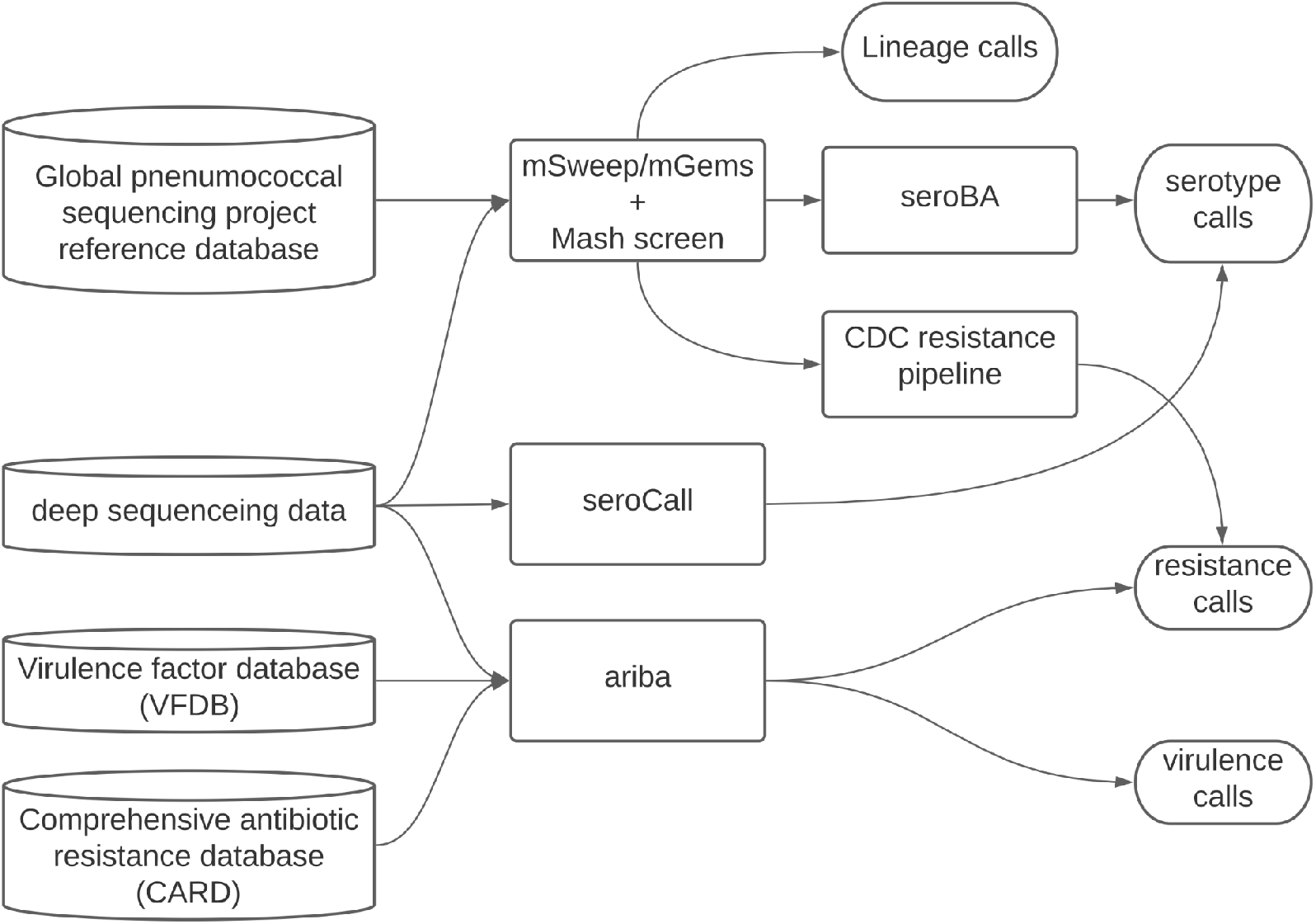
A schematic indicating the bioinformatics pipeline used to call both serotypes and resistance elements from the PDS data.

**Supplementary Figure 11.**
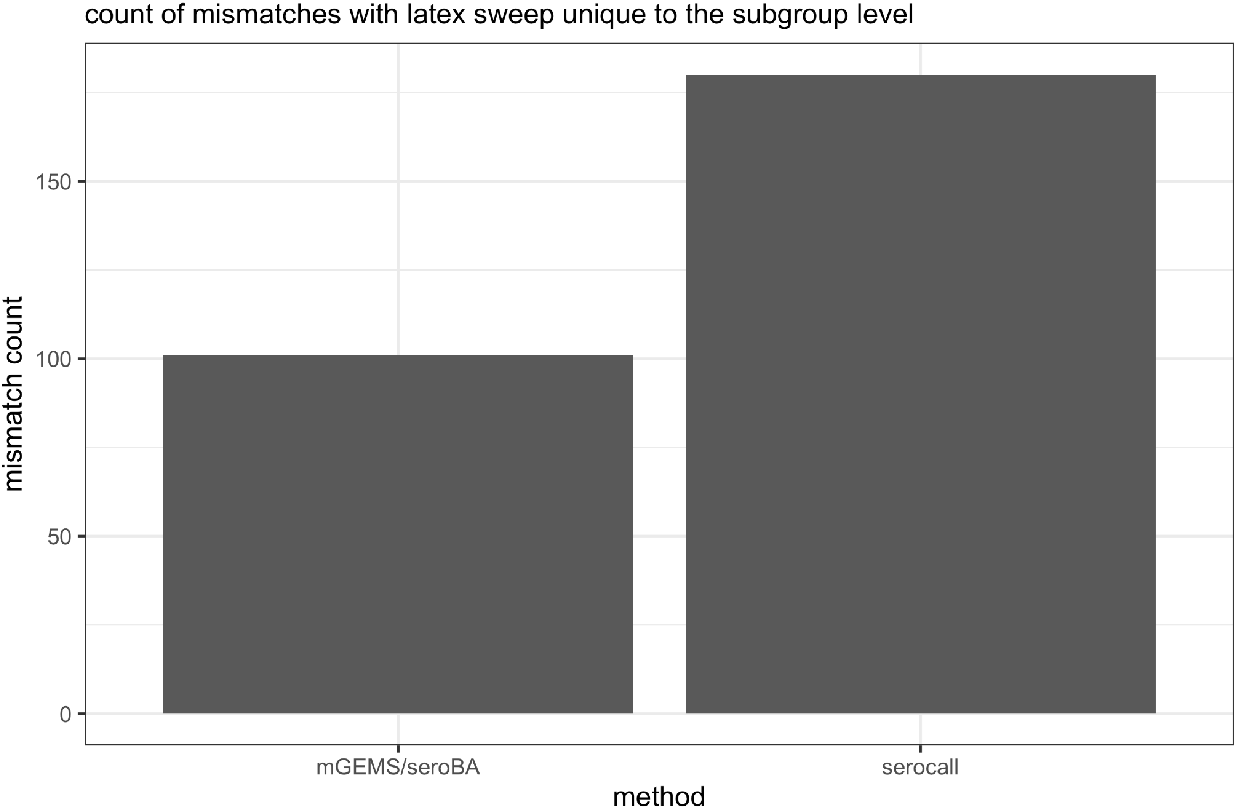
The number of mismatches between latex sweeps and either mGEMS/seroba or the serocall algorithms at the subgroup serotype level. As mGEMs/seroba was found to better agree with latex sweeps in cases of conflict between the two algorithms, the mGEMs/seroba result was chosen.

**Supplementary Figure 12.**
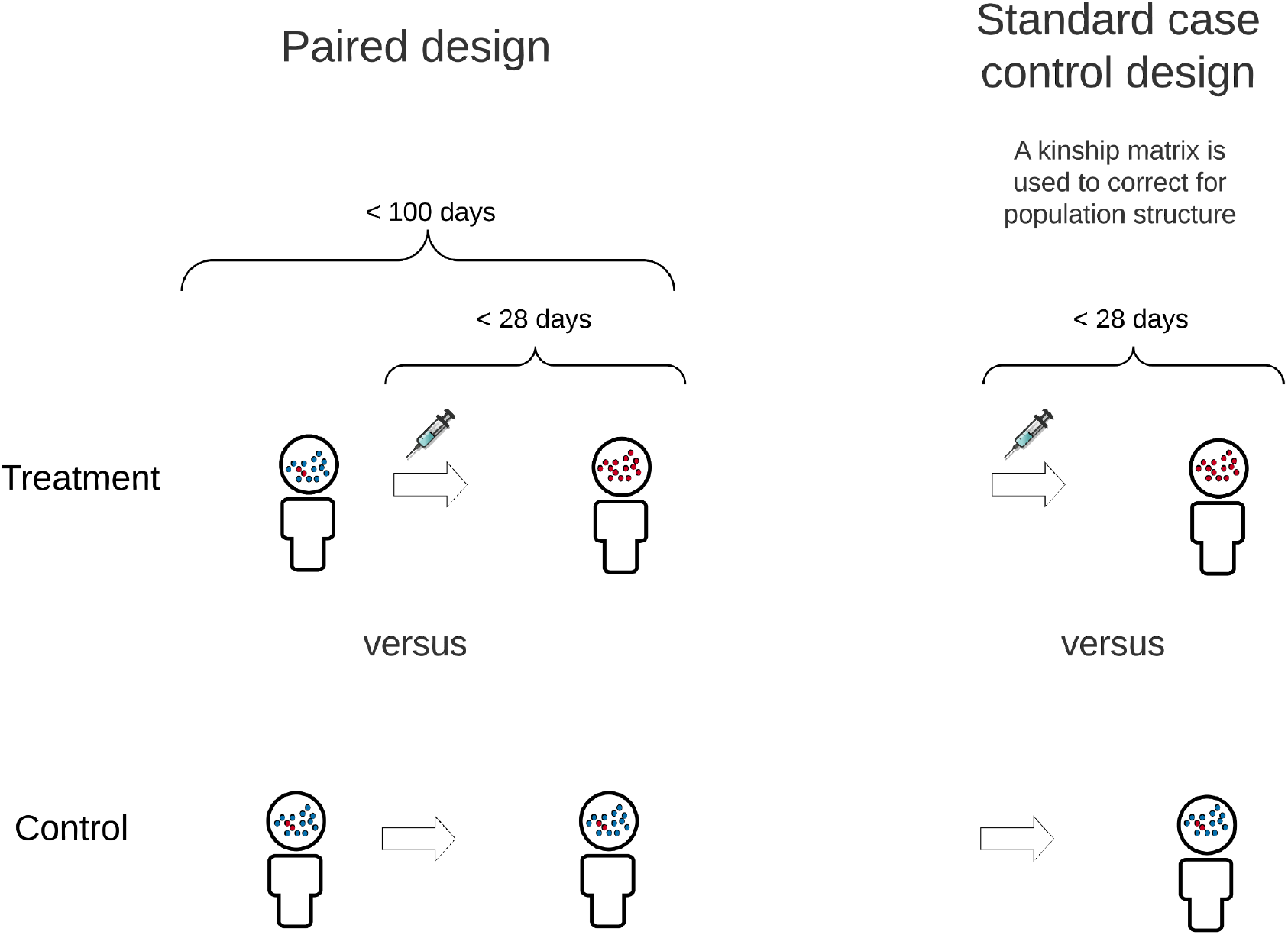
A schematic indicating the design of the two GWAS analyses conducted in this study. A linear model on the log of the unitig counts per million similar to that commonly used in RNA-seq analyses was used in the paired design while the Pyseer algorithm was used in the standard design.

**Supplementary Figure 13.**
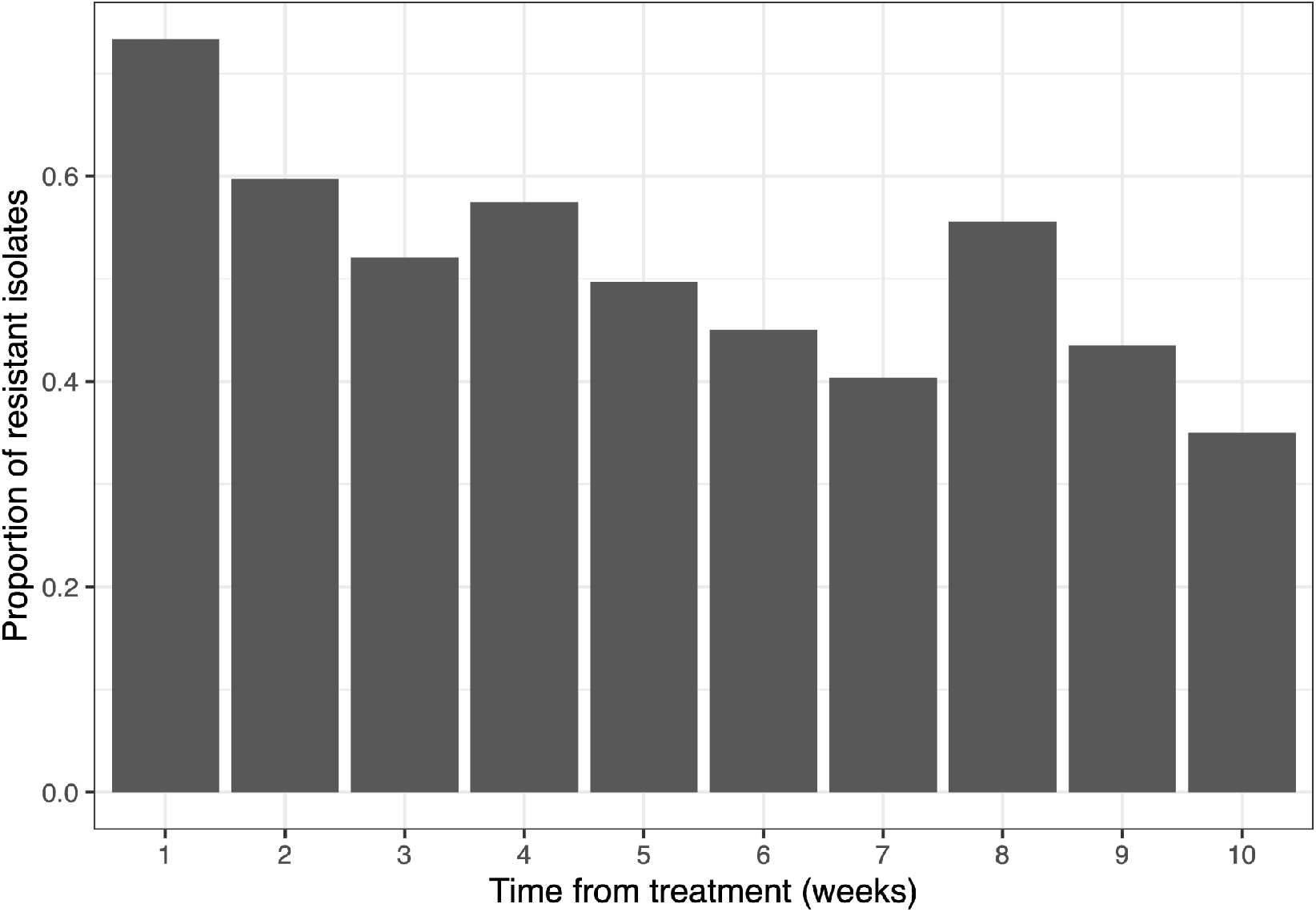
The proportion of resistant isolates following antimicrobial treatment. Those samples within a threshold of 4 weeks (28 days) of a treatment event were classified into the ‘treated’ class.

**Supplementary Figure 14.**
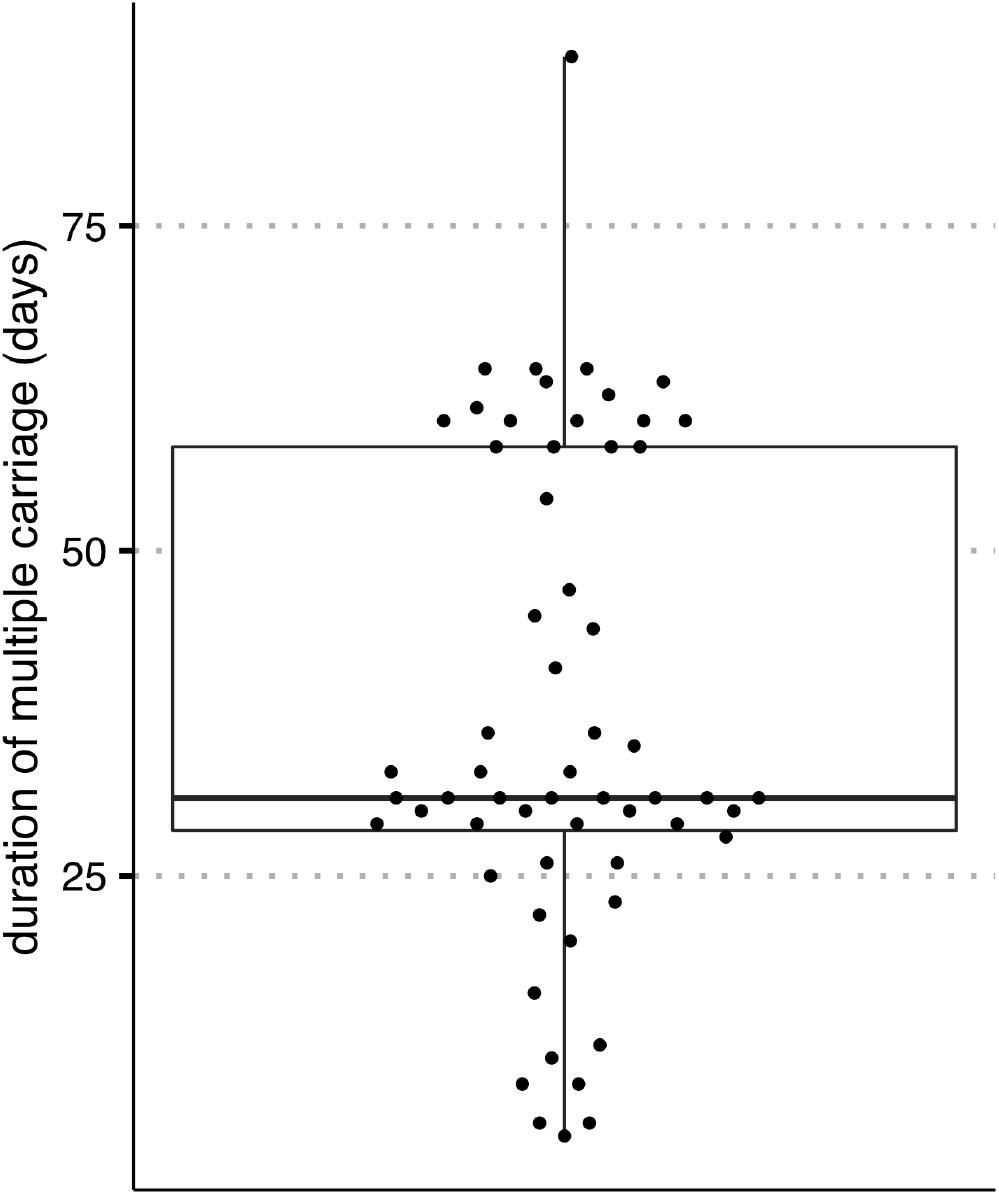
The distribution of the length of time multiple lineages colonised the same host during the same carriage event. Multiple colonisation events that were only observed at a single time point are excluded from this analysis.

**Supplementary Figure 15.**
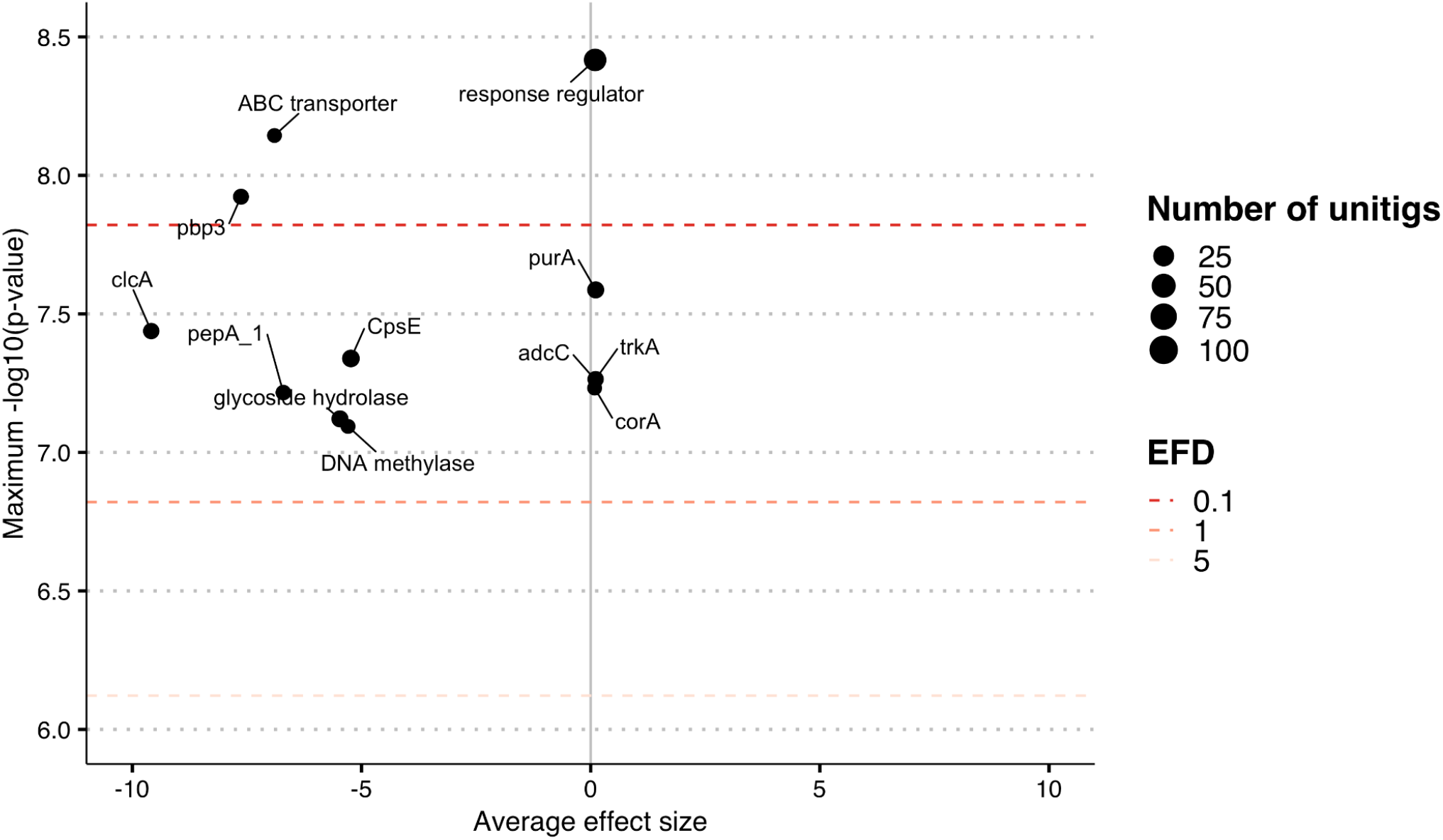
The log odds ratio of clearance following treatment for lineages containing previously classified PBP genes in Li et al., (65) (left). Those in red are significantly more likely to persist following antimicrobial treatment whilst those in blue are likely to be eliminated. The corresponding distribution of MIC profiles for lineages found to contain these PBP genes in Li et al. is given to the right.

**Supplementary Figure 16.**
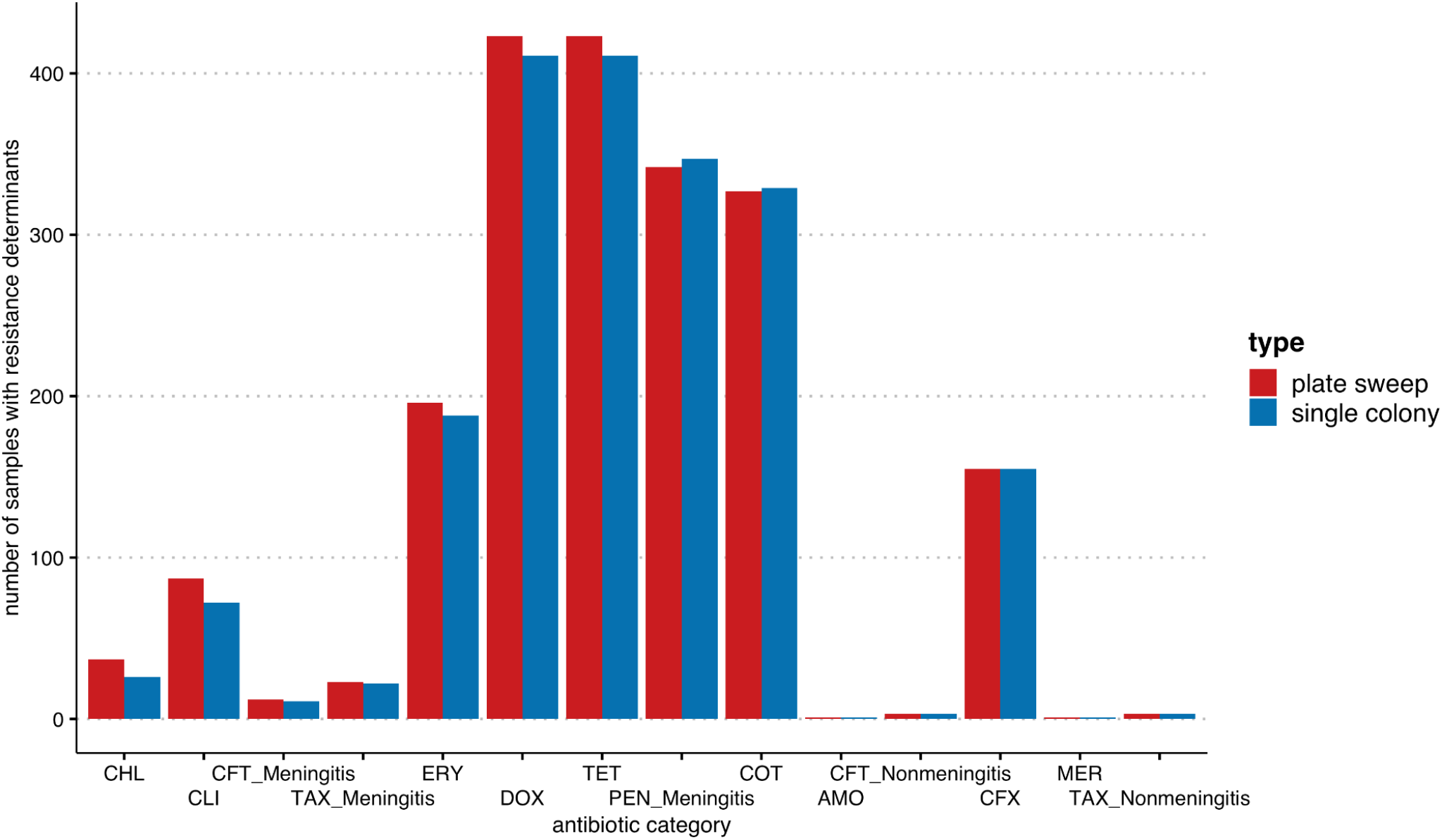
The number of resistance calls identified in 584 samples which consisted of only a single pneumococcal lineage and were sequenced using PDS and via single colony picks in Chewapreecha et al., (24). The high correspondence between the two methods suggests PDS does not lead to a significant number of false positive resistance calls.

**Supplementary Figure 17.**
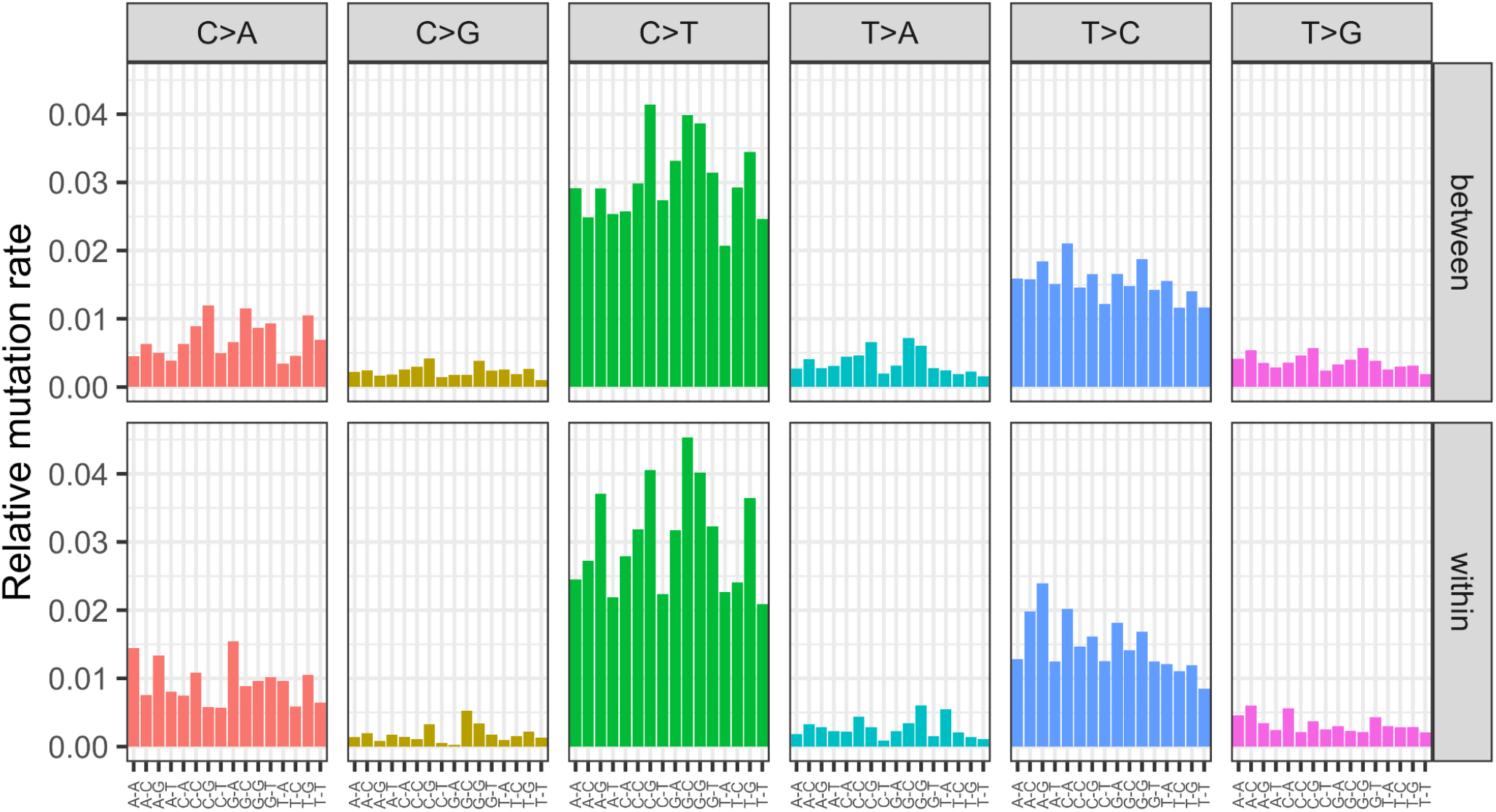
Each spectra is displayed according to the 96 substitution classification defined by the substitution class (colour in the graph) and sequence context immediately 3’ and 5’ to the mutated base. The mutation types are given on the horizontal axes, while vertical axes depict the frequency of each type. Mutations observed using ancestral state reconstruction from genomes observed in different hosts are given above while mutations observed within a host are given in the bottom panels.

**Supplementary Figure 18.**
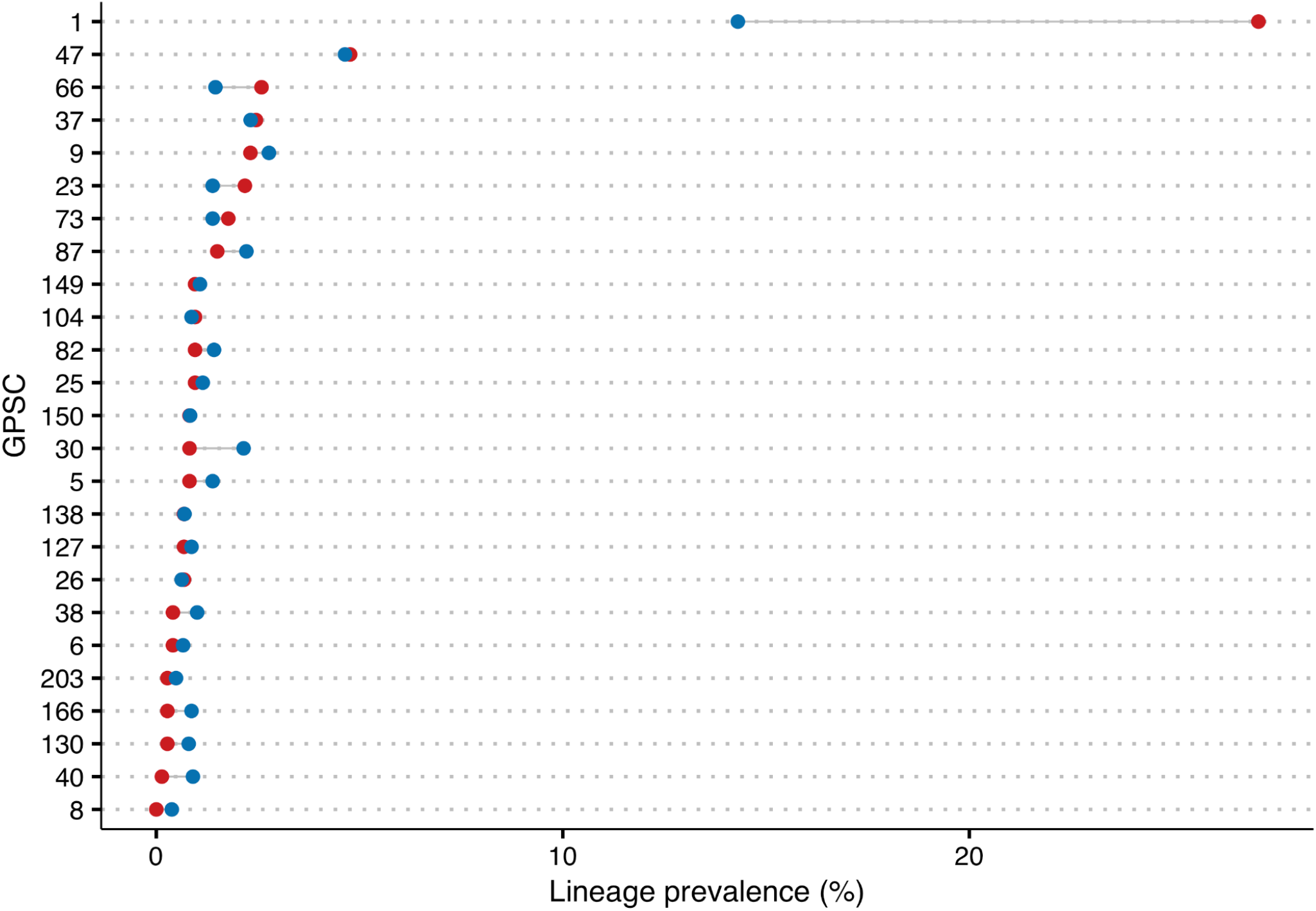
The proportion of lineages made up of each GPSC after treatment (red) and in the absence of treatment (blue). Only those GPSCs that are present at a prevalence of at least 1% in the full data set are included.

**Supplementary Figure 19.**
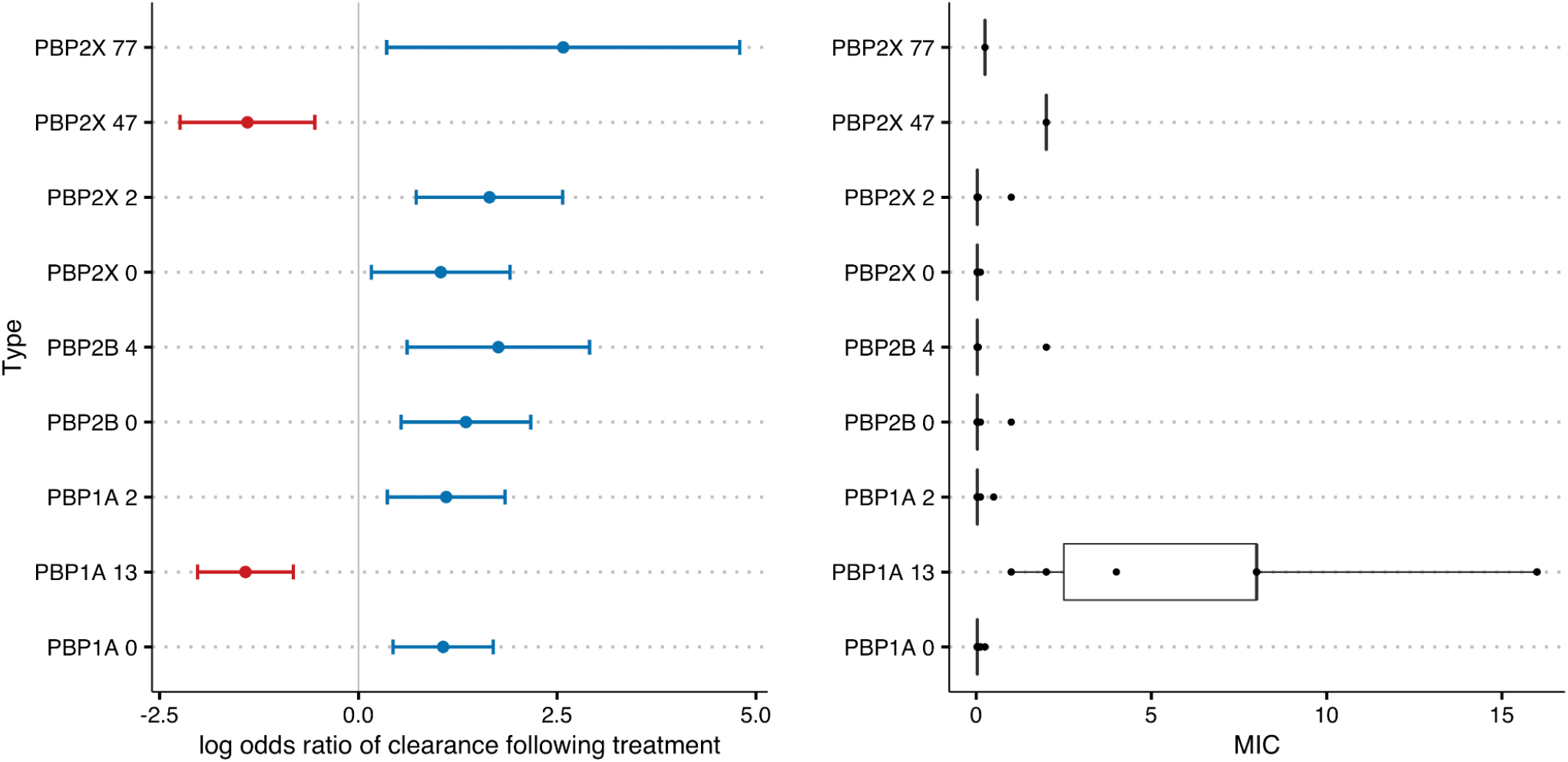
The log odds ratio of clearance following treatment for lineages containing previously classified PBP genes in Li et al., (65) (left). Those in red are significantly more likely to persist following antimicrobial treatment whilst those in blue are likely to be eliminated. The corresponding distribution of MIC profiles for lineages found to contain these PBP genes in Li et al. is given to the right.

## Notes

https://www.ebi.ac.uk/ena/browser/view/PRJEB22771

https://github.com/gtonkinhill/pneumo_withinhost_manuscript

### Competing Interest Statement

The authors have declared no competing interest.

## References

1. Brian Wahl, Katherine L O’Brien, Adena Greenbaum, Anwesha Majumder, Li Liu, Yue Chu, Ivana Lukšić, Harish Nair, David A McAllister, Harry Campbell, Igor Rudan, Robert Black, and Maria Deloria Knoll. Burden of streptococcus pneumoniae and haemophilus influenzae type b disease in children in the era of conjugate vaccines: global, regional, and national estimates for 2000–15. The Lancet Global Health, 6(7):e744–e757, July 2018.

2. GBD 2016 Lower Respiratory Infections Collaborators. Estimates of the global, regional, and national morbidity, mortality, and aetiologies of lower respiratory infections in 195 countries, 1990-2016: a systematic analysis for the global burden of disease study 2016. Lancet Infect. Dis., 18(11):1191–1210, November 2018.

3. Chris Wymant, Matthew Hall, Oliver Ratmann, David Bonsall, Tanya Golubchik, Mariateresa de Cesare, Astrid Gall, Marion Cornelissen, Christophe Fraser, and STOP-HCV Consortium, The Maela Pneumococcal Collaboration, and The BEEHIVE Collaboration. PHYLOSCANNER: Inferring transmission from within- and Between-Host pathogen genetic diversity. Mol. Biol. Evol., November 2017.

4. Finlay Campbell, Camilla Strang, Neil Ferguson, Anne Cori, and Thibaut Jombart. When are pathogen genome sequences informative of transmission events? PLoS Pathog., 14 (2):e1006885, February 2018.

5. Jukka Corander, Christophe Fraser, Michael U Gutmann, Brian Arnold, William P Hanage, Stephen D Bentley, Marc Lipsitch, and Nicholas J Croucher. Frequency-dependent selection in vaccine-associated pneumococcal population dynamics. Nature Ecology & Evolution, page 1, October 2017.

6. Nicholas G Davies, Stefan Flasche, Mark Jit, and Katherine E Atkins. Within-host dynamics shape antibiotic resistance in commensal bacteria. Nature Ecology & Evolution, page 1, February 2019.

7. Taj Azarian, Pamela P Martinez, Brian J Arnold, Xueting Qiu, Lindsay R Grant, Jukka Corander, Christophe Fraser, Nicholas J Croucher, Laura L Hammitt, Raymond Reid, Mathuram Santosham, Robert C Weatherholtz, Stephen D Bentley, Katherine L O’Brien, Marc Lipsitch, and William P Hanage. Frequency-dependent selection can forecast evolution in streptococcus pneumoniae. PLoS Biol., 18(10):e3000878, 2020.

8. Stephanie W Lo, Rebecca A Gladstone, Andries J van Tonder, John A Lees, Mignon du Plessis, Rachel Benisty, Noga Givon-Lavi, Paulina A Hawkins, Jennifer E Cornick, Brenda Kwambana-Adams, Pierra Y Law, Pak Leung Ho, Martin Antonio, Dean B Everett, Ron Dagan, Anne von Gottberg, Keith P Klugman, Lesley McGee, Robert F Breiman, Stephen D Bentley, and Global Pneumococcal Sequencing Consortium. Pneumococcal lineages associated with serotype replacement and antibiotic resistance in childhood invasive pneumococcal disease in the post-PCV13 era: an international whole-genome sequencing study. Lancet Infect. Dis., 19(7):759–769, July 2019.

9. Rebecca A Gladstone, Stephanie W Lo, John A Lees, Nicholas J Croucher, Andries J van Tonder, Jukka Corander, Andrew J Page, Pekka Marttinen, Leon J Bentley, Theresa J Ochoa, Pak Leung Ho, Mignon du Plessis, Jennifer E Cornick, Brenda Kwambana-Adams, Rachel Benisty, Susan A Nzenze, Shabir A Madhi, Paulina A Hawkins, Dean B Everett, Martin Antonio, Ron Dagan, Keith P Klugman, Anne von Gottberg, Lesley McGee, Robert F Breiman, and Stephen D Bentley. International genomic definition of pneumococcal lineages, to contextualise disease, antibiotic resistance and vaccine impact. EBioMedicine, 43:338–346, May 2019.

10. Nicholas J Croucher, Jonathan A Finkelstein, Stephen I Pelton, Patrick K Mitchell, Grace M Lee, Julian Parkhill, Stephen D Bentley, William P Hanage, and Marc Lipsitch. Population genomics of post-vaccine changes in pneumococcal epidemiology. Nat. Genet., 45(6): 656–663, June 2013.

11. Tanya Golubchik, Angela B Brueggemann, Teresa Street, Robert E Gertz, Jr, Chris C A Spencer, Thien Ho, Eleni Giannoulatou, Ruth Link-Gelles, Rosalind M Harding, Bernard Beall, Tim E A Peto, Matthew R Moore, Peter Donnelly, Derrick W Crook, and Rory Bowden. Pneumococcal genome sequencing tracks a vaccine escape variant formed through a multi-fragment recombination event. Nat. Genet., 44(3):352–355, January 2012.

12. L McGee, L McDougal, J Zhou, B G Spratt, F C Tenover, R George, R Hakenbeck, W Hryniewicz, J C Lefévre, A Tomasz, and K P Klugman. Nomenclature of major antimicrobial-resistant clones of streptococcus pneumoniae defined by the pneumococcal molecular epidemiology network. J. Clin. Microbiol., 39(7):2565–2571, July 2001.

13. Arox W Kamng’ona, Jason Hinds, Naor Bar-Zeev, Katherine A Gould, Chrispin Chaguza, Chisomo Msefula, Jennifer E Cornick, Benard W Kulohoma, Katherine Gray, Stephen D Bentley, Neil French, Robert S Heyderman, and Dean B Everett. High multiple carriage and emergence of streptococcus pneumoniae vaccine serotype variants in malawian children. BMC Infect. Dis., 15:234, June 2015.

14. Paul Turner, Jason Hinds, Claudia Turner, Auscharee Jankhot, Katherine Gould, Stephen D Bentley, François Nosten, and David Goldblatt. Improved detection of nasopharyngeal cocolonization by multiple pneumococcal serotypes by use of latex agglutination or molecular serotyping by microarray. J. Clin. Microbiol., 49(5):1784–1789, May 2011.

15. Chrysanti Murad, Eileen M Dunne, Sunaryati Sudigdoadi, Eddy Fadlyana, Rodman Tarigan, Casey L Pell, Emma Watts, Cattram D Nguyen, Catherine Satzke, Jason Hinds, Mia Milanti Dewi, Meita Dhamayanti, Nanan Sekarwana, Kusnandi Rusmil, E Kim Mulholland, and Cissy Kartasasmita. Pneumococcal carriage, density, and co-colonization dynamics: A longitudinal study in indonesian infants. Int. J. Infect. Dis., 86:73–81, September 2019.

16. Philip C Hill, Abiodun Akisanya, Kawsu Sankareh, Yin Bun Cheung, Mark Saaka, George Lahai, Brian M Greenwood, and Richard A Adegbola. Nasopharyngeal carriage of streptococcus pneumoniae in gambian villagers. Clin. Infect. Dis., 43(6):673–679, September 2006.

17. Tanya Golubchik, Elizabeth M Batty, Ruth R Miller, Helen Farr, Bernadette C Young, Hanna Larner-Svensson, Rowena Fung, Heather Godwin, Kyle Knox, Antonina Votintseva, Richard G Everitt, Teresa Street, Madeleine Cule, Camilla L C Ip, Xavier Didelot, Timothy E A Peto, Rosalind M Harding, Daniel J Wilson, Derrick W Crook, and Rory Bowden. Within-host evolution of staphylococcus aureus during asymptomatic carriage. PLoS One, 8(5):e61319, May 2013.

18. Chrispin Chaguza, Madikay Senghore, Ebrima Bojang, Rebecca A Gladstone, Stephanie W Lo, Peggy-Estelle Tientcheu, Rowan E Bancroft, Archibald Worwui, Ebenezer Foster-Nyarko, Fatima Ceesay, Catherine Okoi, Lesley McGee, Keith P Klugman, Robert F Breiman, Michael R Barer, Richard A Adegbola, Martin Antonio, Stephen D Bentley, and Brenda A Kwambana-Adams. Within-host microevolution of streptococcus pneumoniae is rapid and adaptive during natural colonisation. Nat. Commun., 11(1):1–14, July 2020.

19. Xavier Didelot, A Sarah Walker, Tim E Peto, Derrick W Crook, and Daniel J Wilson. Within-host evolution of bacterial pathogens. Nat. Rev. Microbiol., 14(3):150–162, March 2016.

20. J E Barrick and R E Lenski. Genome-wide mutational diversity in an evolving population of escherichia coli. Cold Spring Harb. Symp. Quant. Biol., 74:119–129, January 2009.

21. Robyn S Lee, Jean-François Proulx, Fiona McIntosh, Marcel A Behr, and William P Hanage. Previously undetected super-spreading of mycobacterium tuberculosis revealed by deep sequencing. Elife, 9, February 2020.

22. Josephine M Bryant, Karen P Brown, Sophie Burbaud, Isobel Everall, Juan M Belardinelli, Daniela Rodriguez-Rincon, Dorothy M Grogono, Chelsea M Peterson, Deepshikha Verma, Ieuan E Evans, Christopher Ruis, Aaron Weimann, Divya Arora, Sony Malhotra, Bridget Bannerman, Charlotte Passemar, Kerra Templeton, Gordon MacGregor, Kasim Jiwa, Andrew J Fisher, Tom L Blundell, Diane J Ordway, Mary Jackson, Julian Parkhill, and R Andres Floto. Stepwise pathogenic evolution of mycobacterium abscessus. Science, 372 (6541), April 2021.

23. Tami D Lieberman, Kelly B Flett, Idan Yelin, Thomas R Martin, Alexander J McAdam, Gregory P Priebe, and Roy Kishony. Genetic variation of a bacterial pathogen within individuals with cystic fibrosis provides a record of selective pressures. Nat. Genet., 46(1): 82–87, January 2014.

24. Claire Chewapreecha, Simon R Harris, Nicholas J Croucher, Claudia Turner, Pekka Marttinen, Lu Cheng, Alberto Pessia, David M Aanensen, Alison E Mather, Andrew J Page, Susannah J Salter, David Harris, Francois Nosten, David Goldblatt, Jukka Corander, Julian Parkhill, Paul Turner, and Stephen D Bentley. Dense genomic sampling identifies highways of pneumococcal recombination. Nat. Genet., 46(3):305–309, March 2014.

25. James R Knight, Eileen M Dunne, E Kim Mulholland, Sudipta Saha, Catherine Satzke, Adrienn Tothpal, and Daniel M Weinberger. Determining the serotype composition of mixed samples of pneumococcus using whole-genome sequencing. Microb Genom, 7 (1), January 2021.

26. Sarah Cobey and Marc Lipsitch. Niche and neutral effects of acquired immunity permit coexistence of pneumococcal serotypes. Science, 335(6074):1376–1380, 2012.

27. J A Scott, A J Hall, R Dagan, J M Dixon, S J Eykyn, A Fenoll, M Hortal, L P Jetté, J H Jorgensen, F Lamothe, C Latorre, J T Macfarlane, D M Shlaes, L E Smart, and A Taunay. Serogroup-specific epidemiology of streptococcus pneumoniae: associations with age, sex, and geography in 7,000 episodes of invasive disease. Clin. Infect. Dis., 22(6):973–981, June 1996.

28. Hans-Christian Slotved, Tine Dalby, Zitta Barrella Harboe, Palle Valentiner-Branth, Victoria Fernandez de Casadevante, Laura Espenhain, Kurt Fuursted, and Helle Bossen Konradsen. The incidence of invasive pneumococcal serotype 3 disease in the danish population is not reduced by PCV-13 vaccination. Heliyon, 2(11):e00198, November 2016.

29. Matthias Imöhl, Ralf René Reinert, Christina Ocklenburg, and Mark van der Linden. Association of serotypes of streptococcus pneumoniae with age in invasive pneumococcal disease. J. Clin. Microbiol., 48(4):1291–1296, April 2010.

30. Andreia N Horácio, Catarina Silva-Costa, Joana P Lopes, Mário Ramirez, José Melo-Cristino, Portuguese Group For The Study of Streptococcal Infections, Teresa Vaz, Marília Gião, Rui Ferreira, Ana Buschy Fonseca, Henrique Oliveira, Ana Cristina Silva, Hermínia Costa, Maria Fátima Silva, Maria Amélia Afonso, Margarida Pinto, Odete Chantre, João Marques, Isabel Peres, Isabel Daniel, Ema Canas, Teresa Ferreira, Cristina Marcelo, Lurdes Monteiro, Luís Marques Lito, Filomena Martins, Maria Ana Pessanha, Elsa Gonçalves, Teresa Morais, Teresa Marques, Cristina Toscano, Paulo Lopes, Luísa Felício, Angelina Lameirão, Ana Paula Mota Vieira, Margarida Tomaz, Rosa Bento, Maria Helena Ramos, Castro Ana Paula, Fernando Fonseca, Ana Paula Castro, Graça Ribeiro, Rui Tomé, Celeste Pontes, Luísa Boaventura, Catarina Chaves, Teresa Reis, Nuno Canhoto, Teresa Afonso, Teresa Pina, Helena Peres, Ilse Fontes, Paulo Martinho, Ana Domingos, Gina Marrão, José Grossinho, Manuela Ribeiro, Helena Gonçalves, Alberta Faustino, Adelaide Alves, Maria Cármen Iglesias, Maria Paula Pinheiro, R Semedo, Adriana Coutinho, Luísa Cabral, Olga Neto, Luísa Sancho, José Diogo, Ana Rodrigues, Isabel Nascimento, Elmano Ramalheira, Fernanda Bessa, Raquel Diaz, Isabel Vale, Ana Carvalho, José Miguel Ribeiro, Maria Antónia Read, Valquíria Alves, Margarida Monteiro, Engrácia Raposo, Maria Lurdes Magalhães, Helena Rochas, Anabela Silva, Margarida Rodrigues, José Mota Freitas, Sandra Vieira, Maria Favila Meneses, José Germano de Sousa, Mariana Bettencourt Viana, Isaura Terra, Vitória Rodrigues, Patrícia Pereira, Jesuína Duarte, Paula Pinto, Ezequiel Moreira, João Ataíde Ferreira, Adília Vicente, Paulo Paixão, and Natália Novais. Serotype 3 remains the leading cause of invasive pneumococcal disease in adults in portugal (2012–2014) despite continued reductions in other 13-valent conjugate vaccine serotypes. Front. Microbiol., 7:1616, 2016.

31. Natalie Groves, Carmen L Sheppard, David Litt, Samuel Rose, Ana Silva, Nina Njoku, Sofia Rodrigues, Zahin Amin-Chowdhury, Nicholas Andrews, Shamez Ladhani, and Norman K Fry. Evolution of streptococcus pneumoniae serotype 3 in england and wales: A major vaccine evader. Genes, 10(11), October 2019.

32. Eun Hwa Choi, Fan Zhang, Ying-Jie Lu, and Richard Malley. Capsular polysaccharide (CPS) release by serotype 3 pneumococcal strains reduces the protective effect of AntiType 3 CPS antibodies. Clin. Vaccine Immunol., 23(2):162–167, February 2016.

33. Jo Southern, Nick Andrews, Pamela Sandu, Carmen L Sheppard, Pauline A Waight, Norman K Fry, Albert Jan Van Hoek, and Elizabeth Miller. Pneumococcal carriage in children and their household contacts six years after introduction of the 13-valent pneumococcal conjugate vaccine in england. PLoS One, 13(5):e0195799, May 2018.

34. William P Hausdorff, Daniel R Feikin, and Keith P Klugman. Epidemiological differences among pneumococcal serotypes. Lancet Infect. Dis., 5(2):83–93, February 2005.

35. Neil D Ritchie, Tim J Mitchell, and Tom J Evans. What is different about serotype 1 pneumococci? Future Microbiol., 7(1):33–46, January 2012.

36. Caroline Colijn, Jukka Corander, and Nicholas J Croucher. Designing ecologically optimized pneumococcal vaccines using population genomics. Nat Microbiol, 5(3):473–485, March 2020.

37. Nicola De Maio, Colin J Worby, Daniel J Wilson, and Nicole Stoesser. Bayesian reconstruction of transmission within outbreaks using genomic variants. PLoS Comput. Biol., 14 (4):e1006117, April 2018.

38. James Stimson, Jennifer Gardy, Barun Mathema, Valeriu Crudu, Ted Cohen, and Caroline Colijn. Beyond the SNP threshold: Identifying outbreak clusters using inferred transmissions. Mol. Biol. Evol., 36(3):587–603, March 2019.

39. E Mitsi, A M Roche, J Reiné, T Zangari, J T Owugha, S H Pennington, J F Gritzfeld, A D Wright, A M Collins, S van Selm, M I de Jonge, S B Gordon, J N Weiser, and D M Ferreira. Agglutination by anti-capsular polysaccharide antibody is associated with protection against experimental human pneumococcal carriage. Mucosal Immunol., 10(2):385–394, March 2017.

40. Masamitsu Kono, M Ammar Zafar, Marisol Zuniga, Aoife M Roche, Shigeto Hamaguchi, and Jeffrey N Weiser. Single cell bottlenecks in the pathogenesis of streptococcus pneumoniae. PLoS Pathog., 12(10):e1005887, October 2016.

41. Gerry Tonkin-Hill, Inigo Martincorena, Roberto Amato, Andrew R J Lawson, Moritz Gerstung, Ian Johnston, David K Jackson, Naomi R Park, Stefanie V Lensing, Michael A Quail, Sónia Gonçalves, Cristina Ariani, Michael Spencer Chapman, William L Hamilton, Luke W Meredith, Grant Hall, Aminu S Jahun, Yasmin Chaudhry, Myra Hosmillo, Malte L Pinckert, Iliana Georgana, Anna Yakovleva, Laura G Caller, Sarah L Caddy, Theresa Feltwell, Fahad A Khokhar, Charlotte J Houldcroft, Martin D Curran, Surendra Parmar, Alex Alderton, Rachel Nelson, Ewan Harrison, John Sillitoe, Stephen D Bentley, Jeffrey C Barrett, M Estee Torok, Ian G Goodfellow, Cordelia Langford, Dominic Kwiatkowski, The COVID-19 Genomics UK (COG-UK) Consortium, and Wellcome Sanger Institute COVID-19 Surveillance Team. Patterns of within-host genetic diversity in SARS-CoV-2. December 2020.

42. Matthew D Hall, Matthew Tg Holden, Pramot Srisomang, Weera Mahavanakul, Vanaporn Wuthiekanun, Direk Limmathurotsakul, Kay Fountain, Julian Parkhill, Emma K Nickerson, Sharon J Peacock, and Christophe Fraser. Improved characterisation of MRSA transmission using within-host bacterial sequence diversity. Elife, 8, October 2019.

43. Ellen Heinsbroek, Terence Tafatatha, Christina Chisambo, Amos Phiri, Oddie Mwiba, Bagrey Ngwira, Amelia C Crampin, Jonathan M Read, and Neil French. Pneumococcal acquisition among infants exposed to HIV in rural malawi: A longitudinal household study. Am. J. Epidemiol., 183(1):70–78, January 2016.

44. Tinevimbo Shiri, Kari Auranen, Marta C Nunes, Peter V Adrian, Nadia van Niekerk, Linda de Gouveia, Anne von Gottberg, Keith P Klugman, and Shabir A Madhi. Dynamics of pneumococcal transmission in vaccine-naive children and their HIV-infected or HIV-uninfected mothers during the first 2 years of life. Am. J. Epidemiol., 178(11):1629–1637, December 2013.

45. George Qian, Michiko Toizumi, Sam Clifford, Lien Thuy Le, Tasos Papastylianou, Billy Quilty, Chihiro Iwasaki, Noriko Kitamura, Mizuki Takegata, Trang Minh Nguyen, Hien Anh Thi Nguyen, Duc Anh Dang, Albert Jan van Hoek, Lay Myint Yoshida, and Stefan Flasche. Pneumococcal exposure routes for infants, a nested cross-sectional survey in nha trang, vietnam. medRxiv, page 2021.07.04.21259950, July 2021.

46. Ellen Heinsbroek, Terence Tafatatha, Amos Phiri, Todd D Swarthout, Maaike Alaerts, Amelia C Crampin, Christina Chisambo, Oddie Mwiba, Jonathan M Read, and Neil French. Pneumococcal carriage in households in karonga district, malawi, before and after introduction of 13-valent pneumococcal conjugate vaccination. Vaccine, 36(48):7369–7376, November 2018.

47. Beatriz Maestro and Jesús M Sanz. Choline binding proteins from streptococcus pneumoniae: A dual role as enzybiotics and targets for the design of new antimicrobials. Antibiotics (Basel), 5(2), June 2016.

48. Hilary K DeBardeleben, Elena S Lysenko, Ankur B Dalia, and Jeffrey N Weiser. Tolerance of a phage element by streptococcus pneumoniae leads to a fitness defect during colonization. J. Bacteriol., 196(14):2670–2680, July 2014.

49. Nicholas J Croucher, Joseph J Campo, Timothy Q Le, Xiaowu Liang, Stephen D Bentley, William P Hanage, and Marc Lipsitch. Diverse evolutionary patterns of pneumococcal antigens identified by pangenome-wide immunological screening. Proc. Natl. Acad. Sci. U. S. A., 114(3):E357–E366, January 2017.

50. Peter A Lind and Dan I Andersson. Whole-genome mutational biases in bacteria. Proc. Natl. Acad. Sci. U. S. A., 105(46):17878–17883, November 2008.

51. Justin Jee, Aviram Rasouly, Ilya Shamovsky, Yonatan Akivis, Susan R Steinman, Bud Mishra, and Evgeny Nudler. Rates and mechanisms of bacterial mutagenesis from maximum-depth sequencing. Nature, 534(7609):693–696, June 2016.

52. Prashant Rai, Marcus Parrish, Ian Jun Jie Tay, Na Li, Shelley Ackerman, Fang He, Jimmy Kwang, Vincent T Chow, and Bevin P Engelward. Streptococcus pneumoniae secretes hydrogen peroxide leading to DNA damage and apoptosis in lung cells. Proc. Natl. Acad. Sci. U. S. A., 112(26):E3421–30, June 2015.

53. Iñigo Martincorena, Keiran M Raine, Moritz Gerstung, Kevin J Dawson, Kerstin Haase, Peter Van Loo, Helen Davies, Michael R Stratton, and Peter J Campbell. Universal patterns of selection in cancer and somatic tissues. Cell, 171(5):1029–1041.e21, November 2017.

54. Katherine S Xue and Jesse D Bloom. Linking influenza virus evolution within and between human hosts. Virus Evol, 6(1):veaa010, January 2020.

55. Nicholas J Croucher, Rafal Mostowy, Christopher Wymant, Paul Turner, Stephen D Bentley, and Christophe Fraser. Horizontal DNA transfer mechanisms of bacteria as weapons of intragenomic conflict. PLoS Biol., 14(3):e1002394, March 2016.

56. Nicholas J Croucher, William P Hanage, Simon R Harris, Lesley McGee, Mark van der Linden, Herminia de Lencastre, Raquel Sá-Leão, Jae-Hoon Song, Kwan Soo Ko, Bernard Beall, Keith P Klugman, Julian Parkhill, Alexander Tomasz, Karl G Kristinsson, and Stephen D Bentley. Variable recombination dynamics during the emergence, transmission and ‘disarming’ of a multidrug-resistant pneumococcal clone. BMC Biol., 12:49, June 2014.

57. John A Lees, Nicholas J Croucher, David Goldblatt, François Nosten, Julian Parkhill, Claudia Turner, Paul Turner, and Stephen D Bentley. Genome-wide identification of lineage and locus specific variation associated with pneumococcal carriage duration. Elife, 6, July 2017.

58. Elizabeth M Fozo and Robert G Quivey, Jr. The fabm gene product of streptococcus mutans is responsible for the synthesis of monounsaturated fatty acids and is necessary for survival at low ph. J. Bacteriol., 186(13):4152–4158, July 2004.

59. Konstantinos Papadimitriou, Ángel Alegría, Peter A Bron, Maria de Angelis, Marco Gobbetti, Michiel Kleerebezem, José A Lemos, Daniel M Linares, Paul Ross, Catherine Stanton, Francesca Turroni, Douwe van Sinderen, Pekka Varmanen, Marco Ventura, Manuel Zúñiga, Effie Tsakalidou, and Jan Kok. Stress physiology of lactic acid bacteria. Microbiol. Mol. Biol. Rev., 80(3):837–890, September 2016.

60. Win-Yan Chan, Claire Entwisle, Giuseppe Ercoli, Elise Ramos-Sevillano, Ann McIlgorm, Paola Cecchini, Christopher Bailey, Oliver Lam, Gail Whiting, Nicola Green, David Goldblatt, Jun X Wheeler, and Jeremy S Brown. A novel, Multiple-Antigen pneumococcal vaccine protects against lethal streptococcus pneumoniae challenge. Infect. Immun., 87(3), March 2019.

61. Silvia Altabe, Paloma Lopez, and Diego de Mendoza. Isolation and characterization of unsaturated fatty acid auxotrophs of streptococcus pneumoniae and streptococcus mutans. J. Bacteriol., 189(22):8139–8144, November 2007.

62. M Cyrus Maher, Wondu Alemayehu, Takele Lakew, Bruce D Gaynor, Sara Haug, Vicky Cevallos, Jeremy D Keenan, Thomas M Lietman, and Travis C Porco. The fitness cost of antibiotic resistance in streptococcus pneumoniae: insight from the field. PLoS One, 7(1): e29407, January 2012.

63. Sonja Lehtinen, François Blanquart, Marc Lipsitch, Christophe Fraser, and The Maela Pneumococcal Collaboration. Mechanisms that maintain coexistence of antibiotic sensitivity and resistance also promote high frequencies of multidrug resistance. December 2017.

64. Paul Turner, Claudia Turner, Auscharee Jankhot, Naw Helen, Sue J Lee, Nicholas P Day, Nicholas J White, Francois Nosten, and David Goldblatt. A longitudinal study of streptococcus pneumoniae carriage in a cohort of infants and their mothers on the Thailand-Myanmar border. PLoS One, 7(5):e38271, May 2012.

65. Yuan Li, Benjamin J Metcalf, Sopio Chochua, Zhongya Li, Robert E Gertz, Jr, Hollis Walker, Paulina A Hawkins, Theresa Tran, Cynthia G Whitney, Lesley McGee, and Bernard W Beall. Penicillin-Binding protein transpeptidase signatures for tracking and predicting *β*-Lactam resistance levels in streptococcus pneumoniae. MBio, 7(3), June 2016.

66. E Varon, C Levy, F De La Rocque, M Boucherat, D Deforche, I Podglajen, M Navel, and R Cohen. Impact of antimicrobial therapy on nasopharyngeal carriage of streptococcus pneumoniae, haemophilus influenzae, and branhamella catarrhalis in children with respiratory tract infections. Clin. Infect. Dis., 31(2):477–481, August 2000.

67. Magali Jaillard, Leandro Lima, Maud Tournoud, Pierre Mahé, Alex van Belkum, Vincent Lacroix, and Laurent Jacob. A fast and agnostic method for bacterial genome-wide association studies: Bridging the gap between k-mers and genetic events. PLoS Genet., 14 (11):e1007758, November 2018.

68. John A Lees, Marco Galardini, Stephen D Bentley, Jeffrey N Weiser, and Jukka Corander. pyseer: a comprehensive tool for microbial pangenome-wide association studies. Bioinformatics, 34(24):4310–4312, December 2018.

69. Thierry O Schaffner, Jason Hinds, Katherine A Gould, Daniel Wüthrich, Rémy Bruggmann, Marianne Küffer, Kathrin Mühlemann, Markus Hilty, and Lucy J Hathaway. A point mutation in cpse renders streptococcus pneumoniae nonencapsulated and enhances its growth, adherence and competence. BMC Microbiol., 14:210, August 2014.

70. Mara G Shainheit, Michael D Valentino, Michael S Gilmore, and Andrew Camilli. Mutations in pneumococcal cpse generated via in vitro serial passaging reveal a potential mechanism of reduced encapsulation utilized by a conjunctival isolate. J. Bacteriol., 197(10):1781–1791, May 2015.

71. P David Rogers, Teresa T Liu, Katherine S Barker, George M Hilliard, B Keith English, Justin Thornton, Edwin Swiatlo, and Larry S McDaniel. Gene expression profiling of the response of streptococcus pneumoniae to penicillin. J. Antimicrob. Chemother., 59(4): 616–626, April 2007.

72. Katrina A Lythgoe, Andy Gardner, Oliver G Pybus, and Joe Grove. Short-Sighted virus evolution and a germline hypothesis for chronic viral infections. Trends Microbiol., 25(5): 336–348, May 2017.

73. Claire Chewapreecha, Pekka Marttinen, Nicholas J Croucher, Susannah J Salter, Simon R Harris, Alison E Mather, William P Hanage, David Goldblatt, Francois H Nosten, Claudia Turner, Paul Turner, Stephen D Bentley, and Julian Parkhill. Comprehensive identification of single nucleotide polymorphisms associated with beta-lactam resistance within pneumococcal mosaic genes. PLoS Genet., 10(8):e1004547, August 2014.

74. B G Spratt. Resistance to antibiotics mediated by target alterations. Science, 264(5157): 388–393, April 1994.

75. C G Dowson, T J Coffey, C Kell, and R A Whiley. Evolution of penicillin resistance in streptococcus pneumoniae; the role of streptococcus mitis in the formation of a low affinity PBP2B in s. pneumoniae. Mol. Microbiol., 9(3):635–643, August 1993.

76. Marcin J Skwark, Nicholas J Croucher, Santeri Puranen, Claire Chewapreecha, Maiju Pesonen, Ying Ying Xu, Paul Turner, Simon R Harris, Stephen B Beres, James M Musser, Julian Parkhill, Stephen D Bentley, Erik Aurell, and Jukka Corander. Interacting networks of resistance, virulence and core machinery genes identified by genome-wide epistasis analysis. PLoS Genet., 13(2):e1006508, February 2017.

77. Amilcar J Perez, Michael J Boersma, Kevin E Bruce, Melissa M Lamanna, Sidney L Shaw, Ho-Ching T Tsui, Atsushi Taguchi, Erin E Carlson, Michael S VanNieuwenhze, and Malcolm E Winkler. Organization of peptidoglycan synthesis in nodes and separate rings at different stages of cell division of streptococcus pneumoniae. Mol. Microbiol., 115(6): 1152–1169, June 2021.

78. Nicholas S Briggs, Kevin E Bruce, Souvik Naskar, Malcolm E Winkler, and David I Roper. The pneumococcal divisome: Dynamic control of streptococcus pneumoniae cell division. Front. Microbiol., 12:737396, October 2021.

79. Robert S Brzozowski, Mirella Huber, A Maxwell Burroughs, Gianni Graham, Merryck Walker, Sameeksha S Alva, L Aravind, and Prahathees J Eswara. Deciphering the role of a SLOG superfamily protein YpsA in Gram-Positive bacteria. Front. Microbiol., 10:623, April 2019.

80. Jie Feng, Andréanne Lupien, Hélène Gingras, Jessica Wasserscheid, Ken Dewar, Danielle Légaré, and Marc Ouellette. Genome sequencing of linezolid-resistant streptococcus pneumoniae mutants reveals novel mechanisms of resistance. Genome Res., 19(7):1214–1223, July 2009.

81. J K Morona, A Guidolin, R Morona, D Hansman, and J C Paton. Isolation, characterization, and nucleotide sequence of IS1202, an insertion sequence of streptococcus pneumoniae. J. Bacteriol., 176(14):4437–4443, July 1994.

82. Sine Fjeldhøj, Rikke Pilmann Laursen, Anni Larnkjær, Christian Mølgaard, Kurt Fuursted, Karen Angeliki Krogfelt, and Hans-Christian Slotved. Probiotics and carriage of streptococcus pneumoniae serotypes in danish children, a double-blind randomized controlled trial. Sci. Rep., 8(1):15258, October 2018.

83. Sook-San Wong, Zheng Quan Toh, Eileen M Dunne, E Kim Mulholland, Mimi L K Tang, Roy M Robins-Browne, Paul V Licciardi, and Catherine Satzke. Inhibition of streptococcus pneumoniae adherence to human epithelial cells in vitro by the probiotic lactobacillus rhamnosus GG. BMC Res. Notes, 6:135, April 2013.

84. Katherine L O’brien, Hanna Nohynek, and THE WHO PNEUMOCOCCAL VACCINE TRIALS CARRIAGE WORKING GROUP. Report from a WHO working group: standard method for detecting upper respiratory carriage of streptococcus pneumoniae. Pediatr. Infect. Dis. J., 22(2):e1, February 2003.

85. Paul Turner, Claudia Turner, Auscharee Jankhot, Kawalee Phakaudom, Francois Nosten, and David Goldblatt. Field evaluation of culture plus latex sweep serotyping for detection of multiple pneumococcal serotype colonisation in infants and young children. PLoS One, 8(7):e67933, July 2013.

86. Aarti Desai, Veer Singh Marwah, Akshay Yadav, Vineet Jha, Kishor Dhaygude, Ujwala Bangar, Vivek Kulkarni, and Abhay Jere. Identification of optimum sequencing depth especially for de novo genome assembly of small genomes using next generation sequencing data. PLoS One, 8(4):e60204, April 2013.

87. Tommi Mäklin, Teemu Kallonen, Jarno Alanko, Ørjan Samuelsen, Kristin Hegstad, Veli Mäkinen, Jukka Corander, Eva Heinz, and Antti Honkela. Bacterial genomic epidemiology with mixed samples. Microb Genom, 7(11), November 2021.

88. Tommi Mäklin, Teemu Kallonen, Sophia David, Christine J Boinett, Ben Pascoe, Guillaume Méric, David M Aanensen, Edward J Feil, Stephen Baker, Julian Parkhill, and Others. High-resolution sweep metagenomics using fast probabilistic inference. Wellcome Open Research, 5(14):14, 2020.

89. John A Lees, Simon R Harris, Gerry Tonkin-Hill, Rebecca A Gladstone, Stephanie W Lo, Jeffrey N Weiser, Jukka Corander, Stephen D Bentley, and Nicholas J Croucher. Fast and flexible bacterial genomic epidemiology with PopPUNK. Genome Res., 29(2):304–316, February 2019.

90. Brian D Ondov, Gabriel J Starrett, Anna Sappington, Aleksandra Kostic, Sergey Koren, Christopher B Buck, and Adam M Phillippy. Mash screen: high-throughput sequence containment estimation for genome discovery. Genome Biol., 20(1):232, November 2019.

91. Lennard Epping, Andries J van Tonder, Rebecca A Gladstone, The Global Pneumococcal Sequencing Consortium, Stephen D Bentley, Andrew J Page, and Jacqueline A Keane. SeroBA: rapid high-throughput serotyping of streptococcus pneumoniae from whole genome sequence data. Microb Genom, 4(7), July 2018.

92. B J Metcalf, R E Gertz, Jr, R A Gladstone, H Walker, L K Sherwood, D Jackson, Z Li, C Law, P A Hawkins, S Chochua, M Sheth, N Rayamajhi, S D Bentley, L Kim, C G Whitney, L McGee, B Beall, and Active Bacterial Core surveillance team. Strain features and distributions in pneumococci from children with invasive disease before and after 13-valent conjugate vaccine implementation in the USA. Clin. Microbiol. Infect., 22(1):60.e9–60.e29, January 2016.

93. Brian D Ondov, Todd J Treangen, Páll Melsted, Adam B Mallonee, Nicholas H Bergman, Sergey Koren, and Adam M Phillippy. Mash: fast genome and metagenome distance estimation using MinHash. Genome Biol., 17(1):132, June 2016.

94. John A Lees, Minna Vehkala, Niko Välimäki, Simon R Harris, Claire Chewapreecha, Nicholas J Croucher, Pekka Marttinen, Mark R Davies, Andrew C Steer, Steven Y C Tong, Antti Honkela, Julian Parkhill, Stephen D Bentley, and Jukka Corander. Sequence element enrichment analysis to determine the genetic basis of bacterial phenotypes. Nat. Commun., 7:12797, September 2016.

95. Guillaume Holley and Páll Melsted. Bifrost: highly parallel construction and indexing of colored and compacted de bruijn graphs. Genome Biol., 21(1):249, September 2020.

96. John D Storey and Robert Tibshirani. Statistical significance for genomewide studies. Proc. Natl. Acad. Sci. U. S. A., 100(16):9440–9445, August 2003.

97. Sarah G Earle, Chieh-Hsi Wu, Jane Charlesworth, Nicole Stoesser, N Claire Gordon, Timothy M Walker, Chris C A Spencer, Zamin Iqbal, David A Clifton, Katie L Hopkins, Neil Woodford, E Grace Smith, Nazir Ismail, Martin J Llewelyn, Tim E Peto, Derrick W Crook, Gil McVean, A Sarah Walker, and Daniel J Wilson. Identifying lineage effects when controlling for population structure improves power in bacterial association studies. Nat Microbiol, 1:16041, April 2016.

98. Nathaniel S O’Connell, Lin Dai, Yunyun Jiang, Jaime L Speiser, Ralph Ward, Wei Wei, Rachel Carroll, and Mulugeta Gebregziabher. Methods for analysis of Pre-Post data in clinical research: A comparison of five common methods. J. Biom. Biostat., 8(1):1–8, February 2017.

99. Andreas Wilm, Pauline Poh Kim Aw, Denis Bertrand, Grace Hui Ting Yeo, Swee Hoe Ong, Chang Hua Wong, Chiea Chuen Khor, Rosemary Petric, Martin Lloyd Hibberd, and Niranjan Nagarajan. LoFreq: a sequence-quality aware, ultra-sensitive variant caller for uncovering cell-population heterogeneity from high-throughput sequencing datasets. Nucleic Acids Res., 40(22):11189–11201, December 2012.

100. Stephen J Bush, Dona Foster, David W Eyre, Emily L Clark, Nicola De Maio, Liam P Shaw, Nicole Stoesser, Tim E A Peto, Derrick W Crook, and A Sarah Walker. Genomic diversity affects the accuracy of bacterial single-nucleotide polymorphism-calling pipelines. Gigascience, 9(2), February 2020.

101. Heng Li and Richard Durbin. Fast and accurate short read alignment with Burrows-Wheeler transform. Bioinformatics, 25(14):1754–1760, July 2009.

102. Alistair Miles. pysamstats: A fast python and command-line utility for extracting simple statistics against genome positions based on sequence alignments from a SAM or BAM file.

103. Aaron McKenna, Matthew Hanna, Eric Banks, Andrey Sivachenko, Kristian Cibulskis, Andrew Kernytsky, Kiran Garimella, David Altshuler, Stacey Gabriel, Mark Daly, and Mark A DePristo. The genome analysis toolkit: a MapReduce framework for analyzing next-generation DNA sequencing data. Genome Res., 20(9):1297–1303, September 2010.

104. Robert Dyrdak, Monika Mastafa, Emma B Hodcroft, Richard A Neher, and Jan Albert. Intra- and interpatient evolution of enterovirus D68 analyzed by whole-genome deep sequencing. Virus Evol, 5(1):vez007, January 2019. ISSN 2057-1577. doi: 10.1093/ve/vez007.

105. Nicholas J Croucher, Andrew J Page, Thomas R Connor, Aidan J Delaney, Jacqueline A Keane, Stephen D Bentley, Julian Parkhill, and Simon R Harris. Rapid phylogenetic analysis of large samples of recombinant bacterial whole genome sequences using gubbins. Nucleic Acids Res., page gku1196, November 2014.

106. Yoav Benjamini and Yosef Hochberg. Controlling the false discovery rate: A practical and powerful approach to multiple testing. J. R. Stat. Soc. Series B Stat. Methodol., 57(1): 289–300, January 1995.

107. Bui Quang Minh, Heiko A Schmidt, Olga Chernomor, Dominik Schrempf, Michael D Woodhams, Arndt von Haeseler, and Robert Lanfear. IQ-TREE 2: New models and efficient methods for phylogenetic inference in the genomic era. Mol. Biol. Evol., 37(5):1530–1534, May 2020.

108. N Goldman and Z Yang. A codon-based model of nucleotide substitution for protein-coding DNA sequences. Mol. Biol. Evol., 11(5):725–736, September 1994.

109. Gerry Tonkin-Hill, Neil MacAlasdair, Christopher Ruis, Aaron Weimann, Gal Horesh, John A Lees, Rebecca A Gladstone, Stephanie Lo, Christopher Beaudoin, R Andres Floto, Simon D W Frost, Jukka Corander, Stephen D Bentley, and Julian Parkhill. Producing polished prokaryotic pangenomes with the panaroo pipeline. Genome Biol., 21(1):180, July 2020.

110. Chi C Wong, Inigo Martincorena, Alistair G Rust, Mamunur Rashid, Constantine Alifrangis, Ludmil B Alexandrov, Jessamy C Tiffen, Christina Kober, Chronic Myeloid Disorders Working Group of the International Cancer Genome Consortium, Anthony R Green, Charles E Massie, Jyoti Nangalia, Stella Lempidaki, Hartmut Döhner, Konstanze Döhner, Sarah J Bray, Ultan McDermott, Elli Papaemmanuil, Peter J Campbell, and David J Adams. Inactivating CUX1 mutations promote tumorigenesis. Nat. Genet., 46(1):33–38, January 2014.

111. Eduardo P C Rocha, John Maynard Smith, Laurence D Hurst, Matthew T G Holden, Jessica E Cooper, Noel H Smith, and Edward J Feil. Comparisons of dN/dS are time dependent for closely related bacterial genomes. J. Theor. Biol., 239(2):226–235, March 2006.

112. Sergey Kryazhimskiy and Joshua B Plotkin. The population genetics of dN/dS. PLoS Genet., 4(12):e1000304, December 2008.

113. Thomas P Minka. Estimating a dirichlet distribution. http://citeseerx.ist.psu.edu›viewdoc›summaryhttp://citeseerx.ist.psu.edu›viewdoc›summary, 2000.

114. Gerry Tonkin-Hill. fasttranscluster.

